# Changes of urinary proteome in high-fat diet *ApoE*^-/-^ mice

**DOI:** 10.1101/2022.08.27.505538

**Authors:** Hua Yuanrui, Meng Wenshu, Wei Jing, Liu Yongtao, Gao Youhe

## Abstract

Cardiovascular disease is currently the leading cause of death worldwide. Atherosclerosis is an important pathological basis of cardiovascular disease, and its early diagnosis is of great significance. Urine is more conducive in the accumulation and response of changes in the physiological state of the body and is not regulated by homeostasis mechanisms, so it is a good source of biomarkers in the early stage of disease. In this study, *ApoE*^-/-^ mice were fed with a high-fat diet for 5 months. Urine samples from the experimental group and control group, which were C57BL/6 mice fed a normal diet, were collected at seven time points. Proteomic analysis was used for internalcontrol and intergroup control. Internal control results showed a significant difference in the urinary proteome before and after a 1-week high-fat diet, and several differential proteins have been reported to be associated with atherosclerosis or for use as candidate biomarkers. The results of the intergroup control indicated that the biological process enriched by the GO analysis of the differential proteins corresponded to the progression of atherosclerosis. Differences in chemical modifications of urinary proteins have also been reported to be associated with the disease. This study demonstrates that urinary proteomics has the potential to monitor changes in the body sensitively and provides the possibility of identifying early biomarkers of atherosclerosis.

## 1 Introduction

Atherosclerosis (AS) is the primary pathological basis of cardiovascular disease (CVD)^[1]^, which is the leading cause of death in the world today^[2]^. In 2015, more than 17 million people died of cardiovascular disease, accounting for 31% of all deaths worldwide^[3]^. The impact of atherosclerosis is more significant during the late stages and induces a series of fatal consequences, such as myocardial infarction and stroke. Therefore, its early diagnosis is of vital importance.

Urine is an ideal source of early biomarkers because biomarkers are measurable changes related to biological processes regulated by homeostasis mechanisms, and urine can accumulate these early changes ^[4]^. This conclusion has been confirmed by many related studies. For example, in a glioblastoma animal model constructed by injecting tumour cells into the brains of rats, changes in the urine proteome occurred before magnetic resonance imaging reflected the changes caused by the tumour ^[5]^. Similarly, studies have confirmed that even if only approximately 10 cells are subcutaneously inoculated in rats, the urinary proteome can also change significantly ^[6]^. In addition, urine is more accessible and non-invasive to obtain.

The use of animal models avoids the influence of genetic, environmental and other factors on the urinary proteome, and it is easier to judge the early stage of atherosclerosis and identify biomarkers ^[7]^. Apolipoprotein E (*ApoE*) plays an important role in maintaining the normal levels of cholesterol and triglycerides in serum by transporting lipids in the blood ^[8]^. Mice lacking *ApoE* function develop hypercholesterolemia, increased very-low-density lipoprotein (VLDL) and decreased high-density lipoprotein (HDL), exhibiting spontaneous formation of plaques, and a high-fat diet can greatly accelerate the formation of plaques ^[9]^.

In this experiment, *ApoE*^-/-^ mice were fed with a 5-month high-fat diet. At different timepoints (week 0, week 1, month 1, month 2, month 3, month 4, and month 5) of the experimental process, urine samples were collected and analysed by mass spectrometry. Internal control of the results was conducted as well as intergroup control using urine samples of normal diet C57BL/6 mice. The changes in the proteome and chemical modifications that occur during disease progression provide clues in the search for biomarkers.

## 2 Materials and methods

### 2.1 Experimental animals

Six 4-week-old male *ApoE*^-/-^ mice were purchased from the Laboratory Animal Science Department of Peking University Health Science Centre and fed a high-fat diet (21% fat and 0.15% cholesterol, Beijing Keao Xieli Feed Co., Ltd.) for 5 months. The animal licence is SCXK (Beijing) 2016-0010. A 12-hour normal light-dark cycle and standard temperature (22°C±1°C) and humidity (65%-70%) conditions were used. All animal protocols governing the experiments in this study were approved by the Institute of Basic Medical Sciences Animal Ethics Committee, Peking Union Medical College (Approved ID: ACUC-A02-2014-007). The study was carried out in compliance with the ARRIVE guidelines.

### 2.2 Histopathology

Six 6-month-old *ApoE*^-/-^ mice in the experimental group and four 6-month-old normal diet C57BL/6 mice (purchased from Beijing Vital River Laboratory Animal Technology Co., Ltd.) were euthanized together. Whole arteries were dissected and stained with Oil Red O ^[10]^. The aorta was fixed in 4% paraformaldehyde and dehydrated with isopropanol. After longitudinal incision, it was stained with Oil Red O dye solution (Biotopped, China) for 20 minutes and rinsed three times with isopropanol. A digital camera (Canon, Japan) was then used to obtain images of the aorta, which were analysed using ImageJ software (1.52a, NIH, USA).

### 2.3 Urine collection and sample preparation

To identify the short-term effects of a high-fat diet on animals, urine samples of experimental group mice in week 0 and week 1 were collected during the experiment. To monitor changes in the urinary proteome during the whole process, urine samples of the experimental group at months 1, 2, 3, 4 and 5 were also collected. The urine of four C57BL/6 mice fed a normal diet (all purchased from Beijing Vital River Laboratory Animal Technology Co., Ltd.) corresponding to the age of *ApoE*^-/-^ mice was collected as a control group (not the same batch). All mice were placed in metabolic cages individually for 12 h to collect urine without any treatment. The collected urine samples were immediately stored at −80°C. The experimental process is shown in Figure 1.

**Figure 1.**
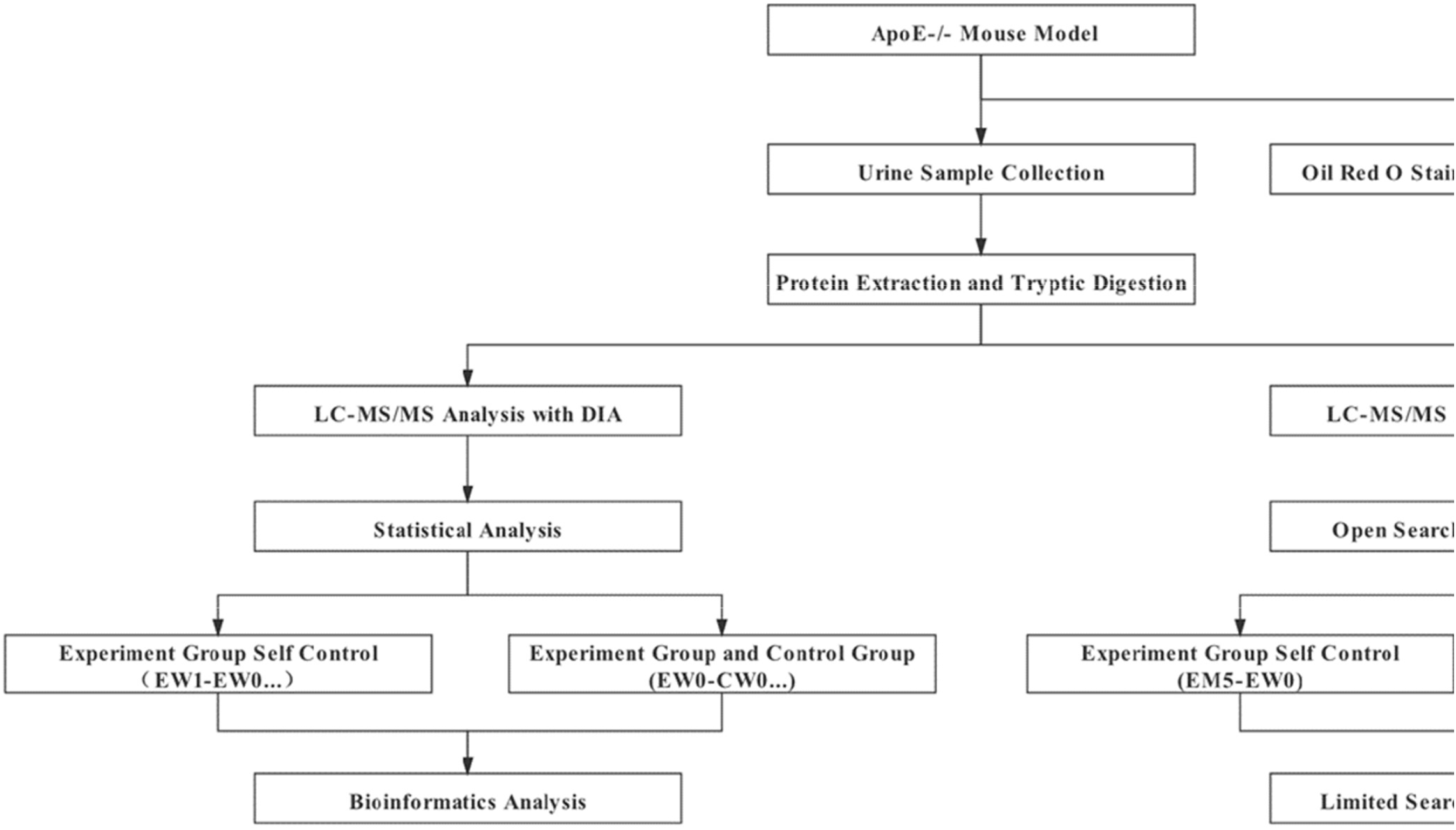
Experimental flow graph

The urine samples were centrifuged at 12,000 g for 40 minutes to remove the supernatant, precipitated using 3 times the volume of ethanol overnight, and then centrifuged at 12,000 g for 30 minutes. The protein was resuspended in lysis buffer (8 mol/L urea, 2 mol/L thiourea, 25 mmol/L dithiothreitol and 50 mmol/L Tris). The protein concentration was measured using the Bradford method. Urine proteolysis was performed using the filter-aided sample preparation (FASP) method^[11]^. The urine protein was loaded on the filter membrane of a 10 kDa ultrafiltration tube (PALL, USA) and washed twice with UA (8 mol/L urea, 0.1 mol/L Tris-HCl, pH 8.5) and 25 mmol/L NH_4_HCO_3_ solution; 20 mmol/L dithiothreitol (DTT, Sigma, USA) was added for reduction at 37°C for 1 hour, and then 50 mmol/L iodoacetamide (IAA, Sigma, USA) was used for alkylation in the dark for 30 minutes. After washing twice with UA and NH_4_HCO_3_ solutions, trypsin (Promega, USA) was added at a ratio of 1:50 for digestion at 37°C for 14 hours. The peptides were passed through Oasis HLB cartridges (Waters, USA) for desalting and then dried by vacuum evaporation (Thermo Fisher Scientific, Germany).

### 2.4 Spin column peptide fractionation

The digested samples were redissolved in 0.1% formic acid and diluted to 0.5 μg/μL. Each sample was used to prepare a mixed peptide sample, and a high pH reversed-phase fractionation spin column (Thermo Fisher Scientific) was used for separation. The mixed peptide samples were added to the chromatographic column and eluted with a step gradient of 8 increasing acetonitrile concentrations (5, 7.5, 10, 12.5, 15, 17.5, 20 and 50% acetonitrile). Ten effluents were finally collected by centrifugation, dried with vacuum evaporation and resuspended in 0.1% formic acid. In this study, iRT reagent (Biognosis, Switzerland) was used to calibrate the retention time of the extracted peptide peaks, which were added to ten components and each sample at a volume ratio of 10:1.

### 2.5 LC-MS/MS analysis

An EASY-nLC 1200 chromatography system (Thermo Fisher Scientific) and Orbitrap Fusion Lumos Tribrid mass spectrometer (Thermo Fisher Scientific) were used for mass spectrometry acquisition and analysis. The peptide sample was loaded onto the precolumn (75 μm×2 cm, C18, 2 μm, Thermo Fisher) at a flow rate of 400 nL/min and then separated using a reversed-phase analysis column (50 μm×15 cm, C18, 2 μm, Thermo Fisher) for 120 minutes. The mobile phase with a gradient of 4%-35% (80% acetonitrile + 0.1% formic acid + 20% water) was used for elution. A full MS scan was acquired within a 350-1,500 m/z range with the resolution set to 120,000. The MS/MS scan was acquired in Orbitrap mode with a resolution of 30,000. The HCD collision energy was set to 30%. The mass spectrum data of 10 components separated by the reversed-phase column and all the samples obtained by enzymatic hydrolysis were collected in DDA mode.

### 2.6 Label-free DIA quantification

The DDA collection results of the above 10 components were imported into the Proteome Discoverer software (version 2.1, Thermo Scientific) search database using the following parameters: mouse database (released in 2019, containing 17038 sequences) with the iRT peptide sequence attached, trypsin digestion, a maximum of two missing cleavage sites are allowed, parent ion mass tolerance is 10 ppm, fragment ion mass tolerance is 0.02 Da, set methionine oxidation as variable modification, set cysteine carbamidomethylation as fixed modification, and protein false discovery rate (FDR) is set to 1%. The PD search result was used to establish the DIA acquisition method and the window width and number were calculated according to the m/z distribution density.

Sixty-nine peptide samples were put into DIA mode to collect mass spectrometry data. Spectronaut™ Pulsar X (Biognosys, Switzerland) software was used to process and analyse mass spectrometry data ^[12]^. According to the DDA search result pdResult file and 10 DDA raw files, create a spectrum library, the raw files collected by DIA were imported for each sample to search the library. The high-confidence protein standard was a peptide q value<0.01, and the peak area of all fragment ions of the secondary peptide was used for protein quantification.

### 2.7 Protein chemical modifications search

PFind Studio software (version 3.1.6, Institute of Computing Technology, Chinese Academy of Sciences) was used to perform label-free quantitative analysis of the DDA collection results of enzymatic hydrolysis samples ^[13]^. The target search database was from the *Mus musculus* database downloaded by UniProt (updated to September 2020). During the search process, the instrument type was set as HCD-FTMS, the enzyme was fully specific trypsin, and up to 2 missed cleaved sites were allowed. The “open-search” mode was selected, and the screening condition was that the FDR at the peptide level was less than 1%. The data used both forward and reverse database search strategies to analyse the data. After the initial screening, a restricted search method was used for verification.

### 2.8 Statistical analysis

The missing abundance values were determined (KNN method) ^[14]^, and CV value screening (CV<0.3) ^[15]^ was performed on the mass spectrometry results. The two-sided unpaired t-test was used for the comparison between each set of data. The internal control of the experimental group and the intergroup control at the same time points were screened for differential proteins. The screening criteria were as follows: fold change (FC) between the two groups ≥1.5 or ≤0.67 and *p <*0.05. At the same time, the samples in each two groups were randomly combined, and the average number of differential proteins in all permutations and combinations was calculated according to the same criteria as normal screening (Table S1), ensuring that differential proteins were generated by differences between groups rather than random production.

The proportions of different types of chemical modification sites in the total modification sites were calculated, and the data between each two groups were compared by two-sided unpaired t-tests. The screening criteria were FC between the two groups ≥1.5 or ≤0.67 and *p <*0.05.

The DAVID database (https://david.ncifcrf.gov) ^[16]^ was used to perform functional enrichment analysis on the differential proteins that were screened. The significance threshold of *p <*0.05 was adopted. All methods were performed in accordance with the relevant guidelines and regulations.

## 3 Results and Discussion

### 3.1 Histopathology

The Oil Red O staining results of the whole aorta of 6-month-old *ApoE*^-/-^ mice fed a high-fat diet for 5 months were compared to those of 6-month-old mice fed a normal diet. The average percentage of stained areas in the experimental group was 17.78±2.14% (n=6), and the average percentage in the control group was 0.88±0.34% (n=4), *p*=0.0004 (Figure 2).

**Figure 2.**
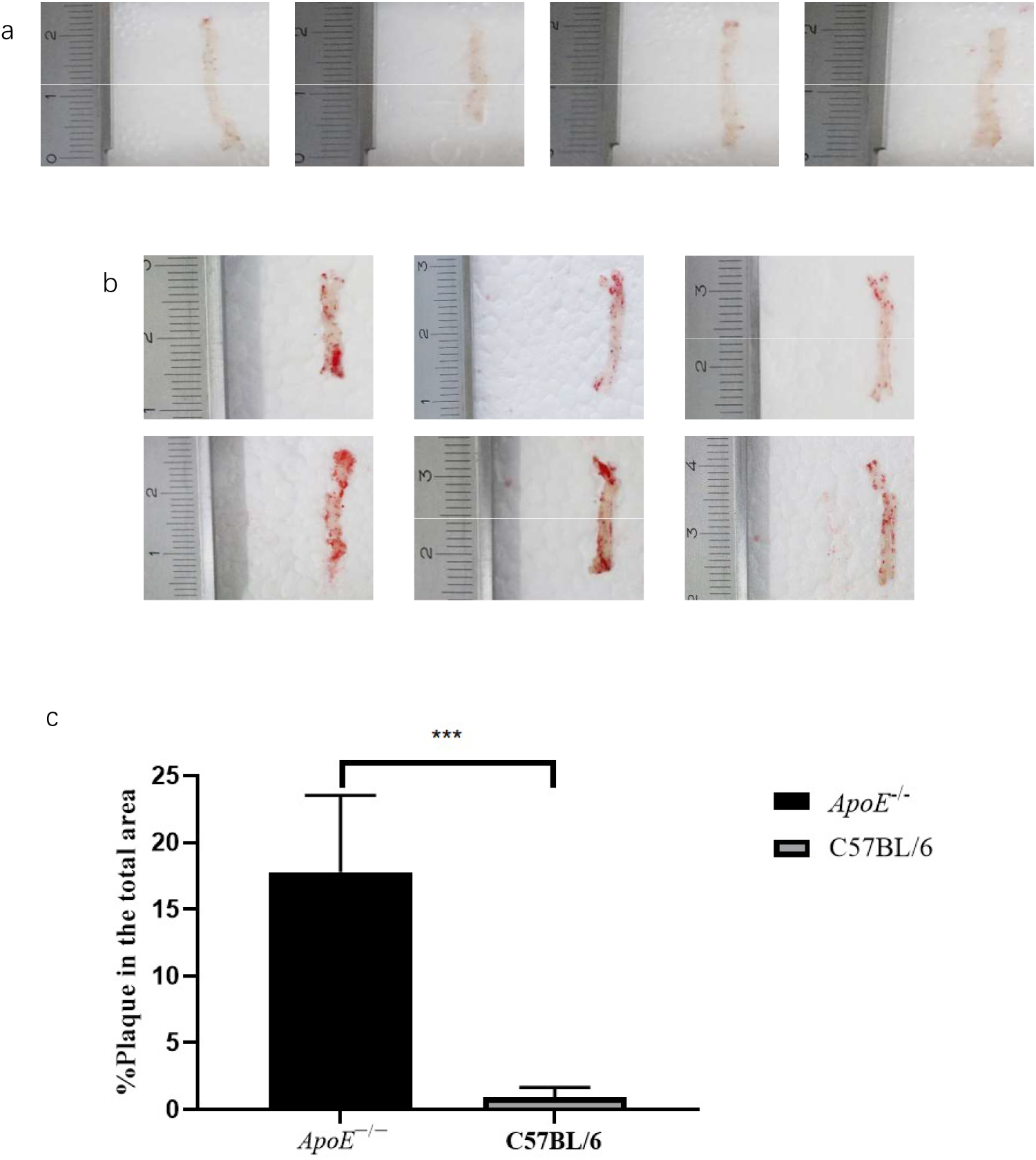
Results and quantitative analysis of oil red O staining of the whole aorta. (a) Results of oil red O staining in the control group; (b) Results of oil red O staining in the experimental group; (c) The comparison of the staining area ratio.

### 3.2 Differential proteins screening and functional annotation

The experimental group and the control group had 69 samples from seven time points (W0/W1/M1/M2/M3/M4/M5) for non-labelled LC-MS/MS quantification (one sample in the experimental group on W0 was insufficient). A total of 592 proteins identified with at least 2 unique peptides with FDR<1% were identified, and an average of 360 urine proteins were identified for each sample. The mass spectrometry proteomics data have been deposited to the ProteomeXchange Consortium (http://proteomecentral.proteomexchange.org) via the iProX partner repository^[17]^ with the dataset identifier <u>PXD027610</u>.

#### 3.2.1 Internal control of the experimental group

##### (1) Short-term effects of a high-fat diet

To identify the effects of a high-fat diet, after a week of a high-fat diet in *ApoE*^-/-^ mice, urine samples collected from W0 and W1 were compared and analysed. A total of 12 proteins were significantly upregulated and 15 proteins were significantly downregulated at W1 (Table 1). Among them, 21 proteins or their family members have been reported to be associated with lipids.

**Table 1.** Details of differential proteins between week 1 and week 0 samples in the experimental group.

| UniProt | Human UniProt | Protein Name | P-value | Fold Change | References |
| --- | --- | --- | --- | --- | --- |
| B5X0G2 | No | Major urinary protein 17 | 0.0008 | 7.92 | [18] |
| P11588 | No | Major urinary protein 1 | 0.0007 | 7.84 | [18] |
| A2BIM8 | No | Major urinary protein 18 | 0.0014 | 3.19 | [18] |
| Q9JI02 | No | Secretoglobin family 2B member 20 | 0.0488 | 2.75 | — |
| Q5FW60 | No | Major urinary protein 20 | 0.0121 | 2.65 | [18] |
| Q07797 | Q08380 | Galectin-3-binding protein | 0.0077 | 2.59 | [60, 61] |
| Q61838 | No | Pregnancy zone protein | 0.0411 | 2.46 | [62] |
| P11591 | No | Major urinary protein 5 | 0.0136 | 2.40 | [18] |
| Q64695 | Q9UNN8 | Endothelial protein C receptor | 0.0150 | 2.23 | [63] |
| Q91WR8 | P59796 | Glutathione peroxidase 6 | 0.0469 | 1.99 | [64] |
| P06797 | P07711 | Cathepsin L1 | 0.0146 | 1.84 | [65] |
| P13597 | P05362 | Intercellular adhesion molecule 1 | 0.0432 | 1.66 | [66, 67] |
| Q9JK39 | A8MVZ5 | Butyrophilin-like protein 10 | 0.0446 | 0.60 | — |
| P01898 | P01891 | H-2 class I histocompatibility antigen, Q10 alpha chain | 0.0429 | 0.59 | [68] |
| P55292 | Q02487 | Desmocollin-2 | 0.0160 | 0.57 | [69, 70] |
| P23780 | P16278 | Beta-galactosidase | 0.0384 | 0.56 | — |
| Q60648 | P17900 | Ganglioside GM2 activator | 0.0382 | 0.56 | [71] |
| P00688 | P04746 | Pancreatic alpha-amylase | 0.0311 | 0.55 | [72] |
| P70269 | P14091 | Cathepsin E | 0.0299 | 0.54 | [73] |
| P11859 | P01019 | Angiotensinogen | 0.0338 | 0.51 | [20] |
| Q6UGQ3 | No | Secretoglobin family 2B member 2 | 0.0270 | 0.49 | — |
| O88322 | Q14112 | Nidogen-2 | 0.0004 | 0.43 | — |
| O88968 | P20062 | Transcobalamin-2 | 0.0155 | 0.39 | [23] |
| P11087 | P02452 | Collagen alpha-1(I) chain | 0.0080 | 0.34 | [21] |
| Q4KML4 | Q9P1F3 | Costars family protein ABRACL | 0.0181 | 0.28 | — |
| P35230 | Q06141 | Regenerating islet-derived protein 3-beta | 0.0012 | 0.24 | [24] |
| A2AEP0 | No | Odorant-binding protein 1b | 0.0213 | 0.20 | [74] |

GO analysis of these 27 proteins by DAVID showed that most of the annotated biological processes were related to lipid metabolism and glucose metabolism (Figure 3). At the same time, the differential proteins between W1 and W0 in the control group (Table S2) did not enrich for any significant changes in biological processes, indicating that the physiological state of mice did not change significantly at W1, while even a week of a high-fat diet induced huge changes in the animal urinary proteome, further demonstrating that the urinary proteome sensitively reflects changes in the body.

**Figure 3.**
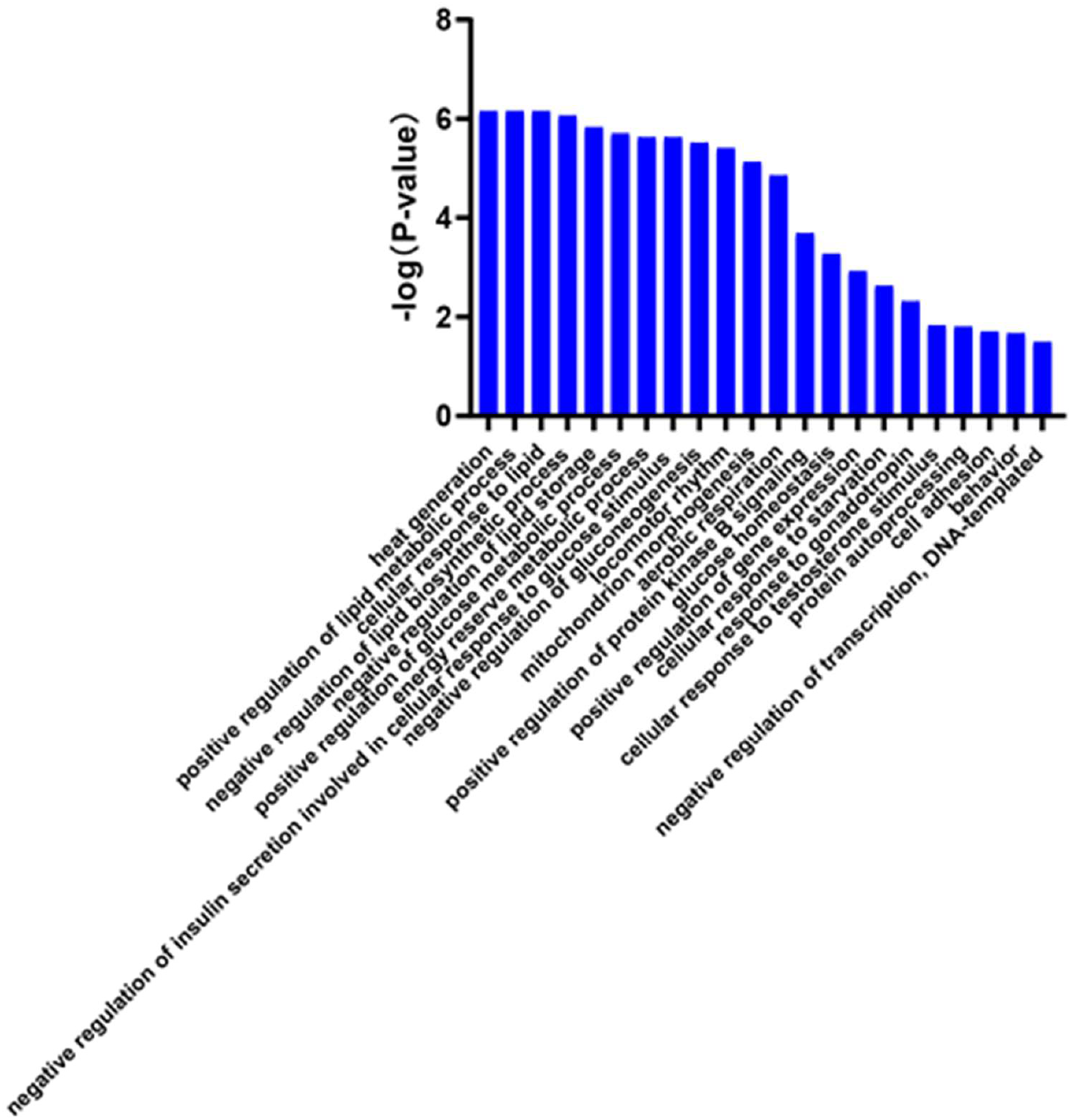
Biological processes enriched in differential proteins between week 1 and week 0 samples of the experimental group (*p <*0.05).

##### (2) Urinary proteome changes in whole process

Compared to W0, 51/69/86/65/88 proteins changed significantly at M1/M2/M3/M4/M5 in the experimental group, respectively. The Venn diagram (Figure 4) shows that a total of 17 proteins changed significantly at all five time points, and the DIA quantitative results show that these 17 proteins exhibited the same change trend at these time points. Another 18 proteins changed significantly at the last four time points, and the trend of change was the same at each time point (Table S3). Among them, 26 proteins or their family members have been previously reported to be related to lipid metabolism or cardiovascular diseases.

**Figure 4.**
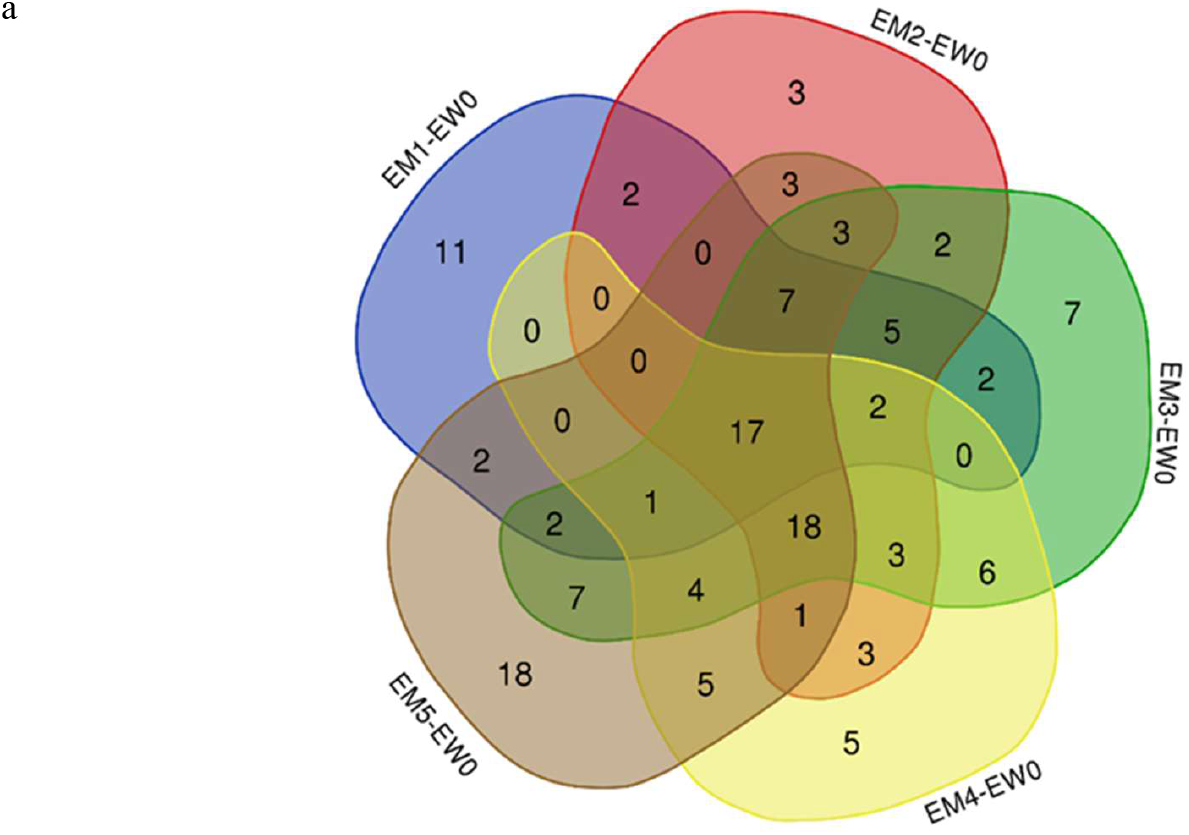
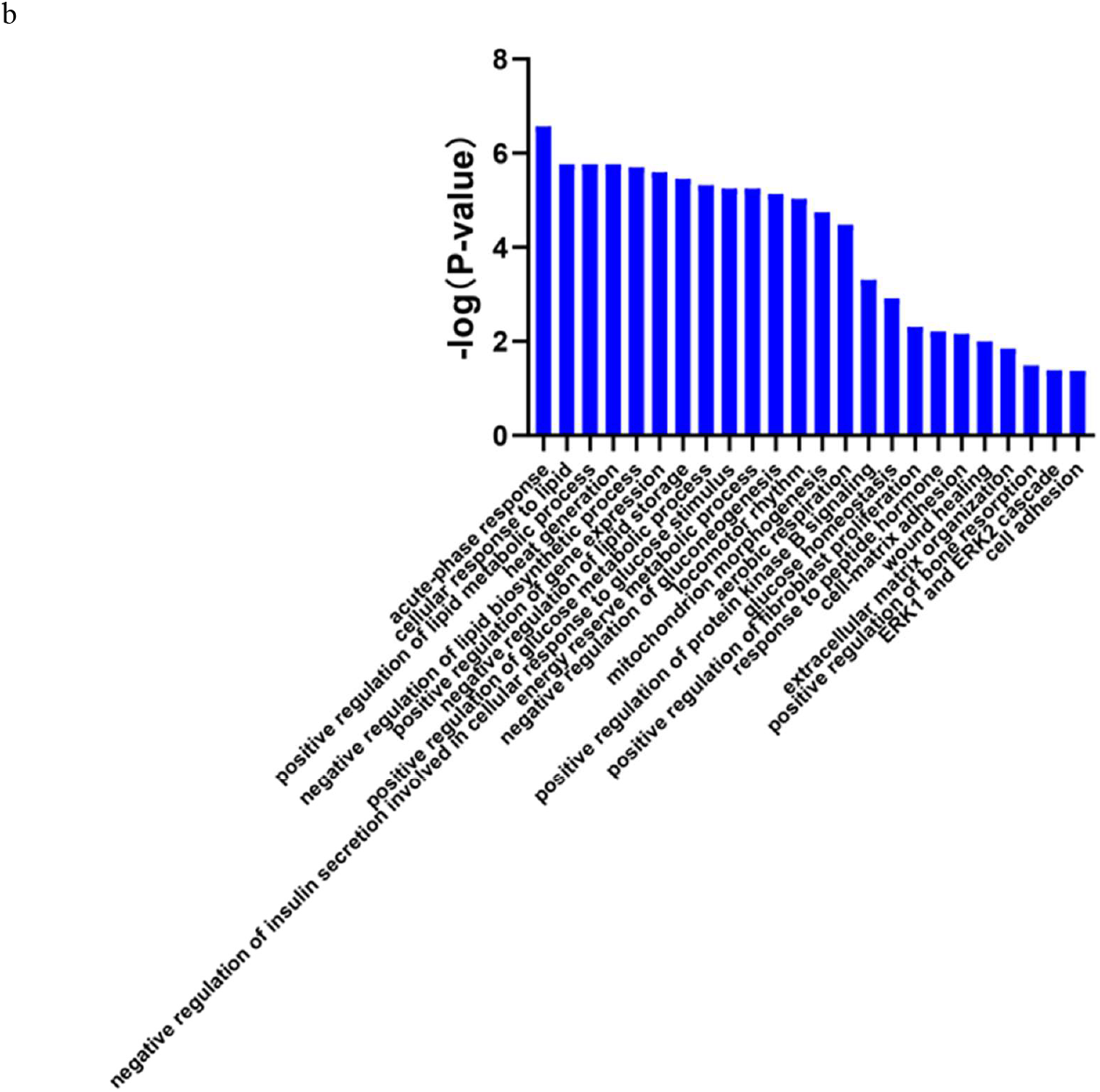
Differential proteins in the whole process. (a) Venn diagram of differential proteins among the other time points (M1/M2/M3/M4/M5) and week 0 samples in the experimental group. (b) Biological processes enriched by continuously changing proteins in the internal control of the experimental group (p <0.05).

Major urinary proteins (MUPs) are members of the lipocalcin family, which can be isolated and transport various lipophilic molecules in the blood and other body fluids ^[18]^. Knockout of mouse trefoil factor 2 protects against obesity in response to a high-fat diet ^[19]^. Angiotensinogen plays a key role in fat cell metabolism and inflammation development ^[20]^. Alpha1-antitrypsin has been reported as a biomarker of atherosclerosis ^[21]^. It has been reported that CCN4 promotes the migration and proliferation of vascular smooth muscle cells by interacting with α5β1 integrin ^[22]^, which plays a vital role in the occurrence and development of atherosclerosis. Regular monitoring of vitamin B12 status may help prevent atherosclerosis-related diseases, and anticobalamin 2 can carry vitamin B12 ^[23]^. Regenerated islet-derived protein 3β, an inflammatory marker, is of great significance for the recruitment of macrophages and tissue repair ^[24]^. The level of α-2-HS-glycoprotein is positively correlated with atherosclerotic substitution parameters, such as intima-media thickness (IMT) and arteriosclerosis ^[25]^. The literature shows that gelsolin stabilizes actin filaments by binding to the ends of filaments, preventing monomer exchange. Its downregulation indicates that the cytoskeleton of vascular smooth muscle cells in the human coronary atherosclerotic medium is dysregulated ^[26]^. It has been reported that *SCUBE2* may play an important role in the progression of atherosclerotic plaques through Hh signal transduction ^[27]^. Type I collagen is an early biomarker of atherosclerosis ^[21]^.

Igκ chain V-III region PC 7043, Igκ chain V-II region 26-10 and immunoglobulin κ constant are all involved in the adaptive immune response. The haptoglobin polymorphism is related to the prevalence and clinical evolution of many inflammatory diseases, including atherosclerosis ^[28]^. MHCII antigen presentation has an important protective function in atherosclerosis ^[29]^. Interleukin-18 plays a key role in atherosclerosis and plays a role in appetite control and the development of obesity ^[30]^. According to the literature, compared to healthy controls, *LAMP-2* gene expression and protein levels in peripheral blood leukocytes of patients with coronary heart disease are significantly increased ^[31]^. T-cadherin is essential for the accumulation of adiponectin in neointima and atherosclerotic plaque lesions ^[32]^. Kidney androgen-regulated protein has also been reported in the urine of *ApoE*^-/-^ mice fed a high-fat diet ^[21]^. Fibronectin is an indicator of connective tissue formation during atherosclerosis ^[33]^. Peripheral arterial occlusive disease (PAOD) is one of the primary manifestations of systemic atherosclerosis, and transthyretin and complement factor B are potential markers for monitoring plasma PAOD disease ^[34]^. Serotransferrin plays an important role in atherosclerosis ^[35]^. The differential expression of serine protease inhibitor A3 in blood vessels is significantly related to human atherosclerosis ^[36]^. Prolactin plays a role in the proliferation of vascular smooth muscle cells, and the proliferation of vascular smooth muscle cells is a characteristic of cardiovascular diseases, such as hypertension and atherosclerosis ^[37]^.

The abovementioned differential proteins that continually changed during the whole process were analysed using DAVID for GO analysis (Figure 4), and the enriched biological processes are also shown in the figure.

The major urinary protein-induced lipid metabolism and glucose metabolism-related biological processes changed significantly; the acute phase reaction has been reported in the literature to be related to atherosclerosis ^[38].^ The positive regulation of fibroblast proliferation also changed significantly, and vascular damage and dysfunction of adipose tissue around blood vessels promotes vasodilation, fibroblast activation and myofibroblast differentiation ^[39]^. Wound healing is also related to atherosclerosis ^[40]^. The extracellular matrix gives atherosclerotic lesion areas tensile strength, viscoelasticity, and compressibility ^[41]^, There are also reports showing a correlation between osteoporosis and atherosclerosis ^[42]^. The ERK1/ERK2 pathway is involved in insulin (INS) and thrombin-induced vascular smooth muscle cells, which play important roles in proliferation ^[43]^. Cell adhesion also plays an important role in atherosclerosis ^[44]^.

The internal control of the experimental group avoids the influence of genetic and dietary differences on the experimental results to the greatest extent, but the influence of biological growth and development is difficult to avoid. The results show that there are a variety of proteins that change continually throughout the progression of the disease and are closely related to the disease. It is worth noting that the differential proteins obtained using this comparison method, and the biological processes and pathways enriched by them exhibited a high degree of overlap at different time points, which may be difficult in enhancing the early diagnosis of the disease, so follow-up intergroup control was performed.

#### 3.2.2 Intergroup control

The comparison of the results between the experimental group and the control group at the same time points showed that 44/16/54/23/48/57/46 differential proteins were obtained at W0/W1/M1/M2/M3/M4/M5, respectively. The details of proteins are shown in Table 2, and the overlap of differential proteins at different time points is shown in Figure S1. Under intergroup control conditions, there were significant differences in the differential proteins at each time point, but they were all closely related to lipids and cardiovascular diseases.

**Table 2.** Details of differential proteins between the experimental group and the control group at different time points.

| UniProt | Human UniProt | Protein Name | Fold Change |  |  |  |  |  |  | References |
| --- | --- | --- | --- | --- | --- | --- | --- | --- | --- | --- |
|  |  |  | EW0-CW0 | EW1-CW1 | EM1-CM1 | EM2-CM2 | EM3-CM3 | EM4-CM4 | EM5-CM5 |  |
| P35230 | Q06141 | Regenerating islet-derived protein 3-beta | 5.22 | — | — | — | — | — | — | [24] |
| P13020 | P06396 | Gelsolin | 2.83 | — | — | — | — | — | — | [26] |
| P97426 | P12724 | Eosinophil cationic protein 1 | 2.57 | — | — | — | 1.77 | 7.57 | — | [75] |
| O88322 | Q14112 | Nidogen-2 | 2.32 | 1.96 | — | — | — | — | 0.39 | — |
| P29699 | P02765 | Alpha-2-HS-glycoprotein | 2.27 | — | — | — | — | — | — | [25] |
| P07758 | P01009 | Alpha-1-antitrypsin 1-1 | 2.11 | — | — | — | 0.58 | 0.25 | — | [21] |
| P07309 | P02766 | Transthyretin | 2.06 | — | — | — | — | — | — | [76] |
| P49183 | P24855 | Deoxyribonuclease-1 | 1.79 | — | 0.47 | — | — | — | — | [77] |
| P01864 | No | Ig gamma-2A chain C region secreted form | 1.55 | — | 0.30 | — | — | — | — | — |
| P19221 | P00734 | Prothrombin | 0.63 | — | — | — | — | — | 0.54 | [78] |
| P61110 | P61109 | Kidney androgen-regulated protein | 0.62 | — | — | — | — | 6.55 | — | [21] |
| Q9Z0M9 | O95998 | Interleukin-18-binding protein | 0.60 | — | — | — | 2.11 | — | — | [30] |
| P05533 | No | Lymphocyte antigen 6A-2/6E-1 | 0.60 | — | — | — | — | — | — | — |
| O09043 | O96009 | Napsin-A | 0.59 | — | — | — | 0.42 | — | — | — |
| P03953 | P00746 | Complement factor D | 0.59 | — | — | — | — | 2.60 | — | [79] |
| Q91VW3 | Q9H299 | SH3 domain-binding glutamic acid-rich-like protein 3 | 0.58 | — | — | 2.22 | — | — | — | — |
| P15379 | P16070 | CD44 antigen | 0.58 | — | 0.25 | — | — | 3.43 | — | [80] |
| P04441 | P04233 | H-2 class II histocompatibility antigen gamma chain | 0.54 | — | — | — | 4.59 | 13.02 | — | [29] |
| P25119 | P20333 | Tumour necrosis factor receptor superfamily member 1B | 0.54 | — | — | — | 1.74 | — | — | [81] |
| Q00993 | P30530 | Tyrosine-protein kinase receptor UFO | 0.53 | — | — | 0.51 | 2.16 | 3.62 | — | [82] |
| P07361 | P02763 | Alpha-1-acid glycoprotein 2 | 0.52 | — | — | — | — | 0.13 | — | [83] |
| P09470 | P12821 | Angiotensin-converting enzyme | 0.51 | — | — | — | — | — | — | [84] |
| Q91WR8 | P59796 | Glutathione peroxidase 6 | 0.47 | — | — | — | — | 3.34 | — | [64] |
| Q62395 | Q07654 | Trefoil factor 3 | 0.46 | — | — | — | — | — | — | [19] |
| Q9DAK9 | Q9NRX4 | 14 kDa phosphohistidine phosphatase | 0.46 | — | — | — | — | — | 2.02 | [85] |
| O88188 | O95711 | Lymphocyte antigen 86 | 0.45 | — | — | — | — | — | — | — |
| Q60932 | P21796 | Voltage-dependent anion-selective channel protein 1 | 0.44 | 0.45 | — | 3.68 | — | — | — | [86] |
| P11589 | No | Major urinary protein 2 | 0.44 | — | 3.82 | — | 1.98 | 6.19 | 3.16 | [18] |
| Q62266 | No | Cornifin-A | 0.43 | 0.49 | — | — | — | — | — | — |
| P17047 | P13473 | Lysosome-associated membrane glycoprotein 2 | 0.42 | — | — | — | 2.19 | 3.45 | 2.44 | [31] |
| P0CW03 | No | Lymphocyte antigen 6C2 | 0.41 | — | — | — | — | — | — | — |
| Q60590 | P02763 | Alpha-1-acid glycoprotein 1 | 0.41 | — | — | — | — | 0.08 | — | [83] |
| P01665 | No | Ig kappa chain V-III region PC 7043 | 0.41 | — | — | — | — | 7.91 | 2.38 | — |
| P04939 | No | Major urinary protein 3 | 0.39 | — | — | — | — | — | — | [18] |
| Q6SJQ5 | Q6UXZ3 | CMRF35-like molecule 3 | 0.35 | — | — | — | — | — | 0.56 | — |
| Q64695 | Q9UNN8 | Endothelial protein C receptor | 0.31 | — | — | — | — | — | — | [63] |
| E9Q557 | P15924 | Desmoplakin | 0.29 | — | — | — | 3.82 | — | — | — |
| P11591 | No | Major urinary protein 5 | 0.27 | — | — | — | — | 4.49 | 2.30 | [18] |
| P51437 | P49913 | Cathelicidin antimicrobial peptide | 0.24 | — | — | — | — | — | — | [87] |
| Q5FW60 | No | Major urinary protein 20 | 0.23 | — | — | — | — | — | 2.14 | [18] |
| A2BIM8 | No | Major urinary protein 18 | 0.20 | 0.50 | 6.10 | — | 2.10 | — | — | [18] |
| Q61646 | P00738 | Haptoglobin | 0.14 | — | — | — | — | 0.09 | — | [28] |
| P11588 | No | Major urinary protein 1 | 0.07 | 0.45 | — | — | 2.17 | — | — | [18] |
| B5X0G2 | No | Major urinary protein 17 | 0.06 | 0.34 | 28.79 | — | — | — | 4.06 | [18] |
| Q9JI02 | No | Secretoglobin family 2B member 20 | — | 2.83 | — | — | 0.23 | — | — | — |
| Q01279 | P00533 | Epidermal growth factor receptor | — | 1.52 | — | — | — | — | — | [88] |
| P10605 | P07858 | Cathepsin B | — | 0.63 | 0.44 | — | — | — | — | [89] |
| P50429 | P15848 | Arylsulfatase B | — | 0.56 | — | — | — | — | — | [90] |
| Q571E4 | P34059 | N-acetylgalactosamine-6-sulfatase | — | 0.52 | — | — | — | 3.57 | — | — |
| Q9JK39 | A8MVZ5 | Butyrophilin-like protein 10 | — | 0.50 | — | — | — | — | — | — |
| P23780 | P16278 | Beta-galactosidase | — | 0.48 | 0.15 | — | 0.28 | — | — | — |
| P70269 | P14091 | Cathepsin E | — | 0.42 | — | — | — | 2.14 | — | [73] |
| O35887 | O43852 | Calumenin | — | 0.40 | — | — | 2.88 | 4.38 | 1.88 | [91] |
| Q9EP95 | Q9BQ08 | Resistin-like alpha | — | 0.27 | — | — | — | — | 0.20 | [92] |
| P20152 | P08670 | Vimentin | — | — | 16.24 | — | — | — | — | [93] |
| Q8K0E8 | P02675 | Fibrinogen beta chain | — | — | 8.28 | — | — | — | — | [94] |
| P16858 | P04406 | Glyceraldehyde-3-phosphate dehydrogenase | — | — | 4.72 | — | — | — | — | [95] |
| Q9WTR5 | P55290 | Cadherin-13 | — | — | 2.78 | — | — | — | — | [32] |
| Q00897 | P01009 | Alpha-1-antitrypsin 1-4 | — | — | 2.57 | — | — | — | 3.27 | [21] |
| O09164 | P08294 | Extracellular superoxide dismutase [Cu-Zn] | — | — | 2.11 | 1.56 | — | 3.37 | — | [96] |
| P11276 | P02751 | Fibronectin | — | — | 0.58 | — | — | 2.15 | — | [33] |
| Q9Z0J0 | P61916 | NPC intracellular cholesterol transporter 2 | — | — | 0.57 | — | — | — | — | [97] |
| Q8BPB5 | Q12805 | EGF-containing fibulin-like extracellular matrix protein 1 | — | — | 0.53 | 0.46 | — | — | — | — |
| Q8BZT5 | Q9H756 | Leucine-rich repeat-containing protein 19 | — | — | 0.52 | 0.50 | 0.55 | — | — | [98] |
| P21614 | P02774 | Vitamin D-binding protein | — | — | 0.50 | — | — | — | 0.49 | [99] |
| P16675 | P10619 | Lysosomal protective protein | — | — | 0.49 | — | — | — | 2.51 | — |
| Q61147 | P00450 | Ceruloplasmin | — | — | 0.49 | — | — | 0.35 | — | [100] |
| P23953 | No | Carboxylesterase 1C | — | — | 0.48 | — | — | — | — | — |
| Q61398 | Q15113 | Procollagen C-endopeptidase enhancer 1 | — | — | 0.48 | — | — | — | — | [101] |
| O35664 | P48551 | Interferon alpha/beta receptor 2 | — | — | 0.47 | — | — | — | — | [102] |
| P11859 | P01019 | Angiotensinogen | — | — | 0.45 | — | — | 0.15 | — | [20] |
| P01898 | P01891 | H-2 class I histocompatibility antigen, Q10 alpha chain | — | — | 0.45 | — | — | — | 2.03 | [68] |
| C0HKG5 | No | Ribonuclease T2-A | — | — | 0.42 | 0.60 | — | — | — | — |
| Q61271 | P36896 | Activin receptor type-1B | — | — | 0.42 | — | — | 3.64 | — | — |
| O88968 | P20062 | Transcobalamin-2 | — | — | 0.40 | — | — | — | — | [23] |
| Q9Z0L8 | Q92820 | Gamma-glutamyl hydrolase | — | — | 0.39 | — | — | — | — | [103, 104] |
| P35459 | Q14210 | Lymphocyte antigen 6D | — | — | 0.39 | 0.57 | — | — | — | — |
| Q61129 | P05156 | Complement factor I | — | — | 0.38 | — | — | — | — | — |
| P01878 | No | Ig alpha chain C region | — | — | 0.38 | — | 5.03 | — | — | — |
| P55292 | Q02487 | Desmocollin-2 | — | — | 0.38 | — | 3.72 | — | — | [69, 70] |
| Q9JJS0 | Q9NQ36 | Signal peptide, CUB and EGF-like domain-containing protein 2 | — | — | 0.35 | — | — | — | — | [27] |
| Q9Z319 | Q9Y5Q5 | Atrial natriuretic peptide-converting enzyme | — | — | 0.35 | — | — | — | — | [105] |
| P09036 | P00995 | Serine protease inhibitor Kazal-type 1 | — | — | 0.34 | — | — | — | — | — |
| Q4KML4 | Q9P1F3 | Costars family protein ABRACL | — | — | 0.33 | — | 2.18 | — | — | — |
| Q925F2 | Q96AP7 | Endothelial cell-selective adhesion molecule | — | — | 0.32 | — | — | — | 0.48 | [106] |
| O89020 | P43652 | Afamin | — | — | 0.31 | 0.50 | — | — | — | [107] |
| Q9DAU7 | Q14508 | WAP four-disulfide core domain protein 2 | — | — | 0.31 | — | — | — | — | — |
| Q8BND5 | O00391 | Sulfhydryl oxidase 1 | — | — | 0.30 | 0.40 | — | — | 0.61 | — |
| P09803 | P12830 | Cadherin-1 | — | — | 0.29 | — | 2.78 | 2.34 | — | [28] |
| P02816 | P12273 | Prolactin-inducible protein homolog | — | — | 0.27 | — | 0.45 | — | — | [37] |
| Q91WR6 | Q9NU53 | Glycoprotein integral membrane protein 1 | — | — | 0.26 | 0.59 | — | — | 0.45 | — |
| Q3UDR8 | Q9GZM5 | Protein YIPF3 | — | — | 0.26 | — | — | — | — | — |
| Q6UGQ3 | No | Secretoglobin family 2B member 2 | — | — | 0.25 | — | — | — | 2.03 | — |
| Q9D3H2 | No | Odorant-binding protein 1a | — | — | 0.25 | — | 1.75 | — | — | [74] |
| P20060 | P07686 | Beta-hexosaminidase subunit beta | — | — | 0.23 | — | 0.35 | — | — | — |
| Q8K1H9 | Q9NY56 | Odorant-binding protein 2a | — | — | 0.21 | — | 0.31 | 0.21 | — | [74] |
| A2AEP0 | No | Odorant-binding protein 1b | — | — | 0.20 | — | — | — | — | [74] |
| Q8C6C9 | Q6P5S2 | Protein LEG1 homolog | — | — | 0.15 | — | 0.44 | — | — | — |
| P00688 | P04746 | Pancreatic alpha-amylase | — | — | 0.14 | — | — | 0.32 | — | [72] |
| P10287 | P22223 | Cadherin-3 | — | — | 0.13 | — | 3.19 | 5.62 | — | [28] |
| P56386 | P60022 | Beta-defensin 1 | — | — | — | 3.94 | — | 5.25 | 1.90 | [108] |
| P10639 | P10599 | Thioredoxin | — | — | — | 3.63 | — | — | — | [109] |
| O88844 | O75874 | Isocitrate dehydrogenase [NADP] cytoplasmic | — | — | — | 2.71 | — | — | — | [110] |
| Q00623 | P02647 | Apolipoprotein A-I | — | — | — | 2.07 | — | — | 0.36 | [111] |
| P0CG49 | P0CG47 | Polyubiquitin-B | — | — | — | 1.98 | — | — | — | [112] |
| Q03404 | Q03403 | Trefoil factor 2 | — | — | — | 1.69 | 2.11 | 8.94 | — | [19] |
| P15947 | P06870 | Kallikrein-1 | — | — | — | 0.66 | — | 1.74 | — | [113] |
| O55186 | P13987 | CD59A glycoprotein | — | — | — | 0.61 | — | 2.53 | — | [114] |
| Q921I1 | P02787 | Serotransferrin | — | — | — | 0.57 | 0.43 | 0.09 | 0.21 | [35] |
| Q60648 | P17900 | Ganglioside GM2 activator | — | — | — | 0.51 | — | — | 2.27 | [72] |
| O88792 | Q9Y624 | Junctional adhesion molecule A | — | — | — | 0.49 | — | — | — | [115] |
| P07724 | P02768 | Albumin | — | — | — | 0.38 | — | 0.38 | 0.43 | [116] |
| O70554 | No | Small proline-rich protein 2B | — | — | — | — | 7.98 | — | 3.26 | [117, 118] |
| Q62267 | No | Cornifin-B | — | — | — | — | 5.69 | — | — | — |
| P35700 | Q06830 | Peroxiredoxin-1 | — | — | — | — | 3.75 | — | — | [119] |
| P18761 | P23280 | Carbonic anhydrase 6 | — | — | — | — | 3.37 | — | — | [120] |
| P01631 | No | Ig kappa chain V-II region 26-10 | — | — | — | — | 3.29 | 3.00 | — | — |
| P11087 | P02452 | Collagen alpha-1(I) chain | — | — | — | — | 3.21 | — | — | [21] |
| P01837 | P01834 | Immunoglobulin kappa constant | — | — | — | — | 3.19 | — | — | — |
| Q99N23 | No | Carbonic anhydrase 15 | — | — | — | — | 2.91 | — | — | [120] |
| P10126 | P68104 | Elongation factor 1-alpha 1 | — | — | — | — | 2.90 | — | — | [121] |
| P70663 | Q14515 | SPARC-like protein 1 | — | — | — | — | 2.74 | — | — | — |
| Q60847 | Q99715 | Collagen alpha-1(XII) chain | — | — | — | — | 2.68 | — | — | [21] |
| P29533 | P19320 | Vascular cell adhesion protein 1 | — | — | — | — | 2.62 | 2.12 | — | [122] |
| O55135 | P56537 | Eukaryotic translation initiation factor 6 | — | — | — | — | 2.55 | — | — | — |
| O54775 | O95388 | CCN family member 4 | — | — | — | — | 2.47 | — | 0.47 | [22] |
| P01843 | No | Ig lambda-1 chain C region | — | — | — | — | 2.45 | — | — | — |
| Q9DBV4 | Q9BRK3 | Matrix remodeling-associated protein 8 | — | — | — | — | 1.70 | — | 1.80 | — |
| O35608 | O15123 | Angiopoietin-2 | — | — | — | — | 1.61 | — | — | [123] |
| Q07797 | Q08380 | Galectin-3-binding protein | — | — | — | — | 1.59 | — | — | [60] |
| Q04519 | P17405 | Sphingomyelin phosphodiesterase | — | — | — | — | 0.57 | — | 1.89 | [124] |
| Q61838 | No | Pregnancy zone protein | — | — | — | — | 0.52 | — | — | [62] |
| Q08423 | P04155 | Trefoil factor 1 | — | — | — | — | — | 12.79 | — | [19] |
| O08997 | O00244 | Copper transport protein ATOX1 | — | — | — | — | — | 8.07 | — | [125] |
| P03977 | No | Ig kappa chain V-III region 50S10.1 | — | — | — | — | — | 4.79 | — | — |
| P42567 | P42566 | Epidermal growth factor receptor substrate 15 | — | — | — | — | — | 4.53 | — | [88] |
| Q91X17 | P07911 | Uromodulin | — | — | — | — | — | 4.22 | — | [126] |
| P57096 | O43653 | Prostate stem cell antigen | — | — | — | — | — | 3.53 | — | — |
| P70699 | P10253 | Lysosomal alpha-glucosidase | — | — | — | — | — | 3.37 | — | [127] |
| P01132 | P01133 | Pro-epidermal growth factor | — | — | — | — | — | 3.13 | — | — |
| P32507 | Q92692 | Nectin-2 | — | — | — | — | — | 3.03 | — | [128] |
| Q5SSE9 | Q86UQ4 | ATP-binding cassette sub-family A member 13 | — | — | — | — | — | 2.92 | — | — |
| P06797 | O60911 | Cathepsin L1 | — | — | — | — | — | 2.63 | — | [65, 73] |
| Q8R242 | Q01459 | Di-N-acetylchitobiase | — | — | — | — | — | 2.47 | — | — |
| Q9Z0K8 | O95497 | Pantetheinase | — | — | — | — | — | 2.32 | 3.65 | — |
| O54782 | Q9Y2E5 | Epididymis-specific alpha-mannosidase | — | — | — | — | — | 1.99 | — | — |
| P22599 | P01009 | Alpha-1-antitrypsin 1-2 | — | — | — | — | — | 0.50 | — | [21] |
| P13634 | P00915 | Carbonic anhydrase 1 | — | — | — | — | — | 0.39 | 3.37 | [120] |
| P20918 | P00747 | Plasminogen | — | — | — | — | — | 0.32 | 0.63 | [129] |
| P07759 | P01011 | Serine protease inhibitor A3K | — | — | — | — | — | 0.26 | — | [36] |
| P02088 | P68871 | Hemoglobin subunit beta-1 | — | — | — | — | — | 0.24 | — | — |
| P01027 | P01024 | Complement C3 | — | — | — | — | — | 0.20 | — | [54] |
| Q00898 | P01009 | Alpha-1-antitrypsin 1-5 | — | — | — | — | — | 0.18 | — | [21] |
| P11672 | P80188 | Neutrophil gelatinase-associated lipocalin | — | — | — | — | — | 0.04 | — | [130] |
| P01887 | P61769 | Beta-2-microglobulin | — | — | — | — | — | — | 3.07 | [131] |
| O09114 | P41222 | Prostaglandin-H2 D-isomerase | — | — | — | — | — | — | 2.41 | [132] |
| Q9WUU7 | Q9UBR2 | Cathepsin Z | — | — | — | — | — | — | 2.32 | [133] |
| Q8BHC0 | Q9Y5Y7 | Lymphatic vessel endothelial hyaluronic acid receptor 1 | — | — | — | — | — | — | 0.61 | — |
| O70570 | P01833 | Polymeric immunoglobulin receptor | — | — | — | — | — | — | 0.55 | — |
| P26041 | P26038 | Moesin | — | — | — | — | — | — | 0.53 | [134] |
| Q7TMJ8 | Q96FE7 | Phosphoinositide-3-kinase-interacting protein 1 | — | — | — | — | — | — | 0.52 | [135] |
| P01660 | No | Ig kappa chain V-III region PC 3741/TEPC 111 | — | — | — | — | — | — | 0.43 | — |
| Q60928 | P19440 | Glutathione hydrolase 1 proenzyme | — | — | — | — | — | — | 0.39 | [104] |
| P61971 | P61970 | Nuclear transport factor 2 | — | — | — | — | — | — | 0.36 | — |
| P06330 | No | Ig heavy chain V region AC38 205.12 | — | — | — | — | — | — | 0.36 | — |
| Q921W8 | Q8WVN6 | Secreted and transmembrane protein 1A | — | — | — | — | — | — | 0.35 | — |
| Q8BX43 | Q969Z4 | Tumour necrosis factor receptor superfamily member 19L | — | — | — | — | — | — | 0.33 | [81] |
| Q9JM99 | Q92954 | Proteoglycan 4 | — | — | — | — | — | — | 0.26 | [135] |

The differential proteins were analysed by DAVID for GO analysis, and the biological processes that changed significantly at different time points are shown in Figure 5. The biological processes related to lipid metabolism and glucose metabolism in the experimental group were significantly different from those in the control group at W0. At W0 and M4, the immune-related processes were significantly different. Differential proteins at M1 were primarily enriched in cell adhesion-related processes, while at M2, they were primarily enriched in redox reaction-related processes. At M3, wound healing began to appear, and there were many adhesion-related processes. In addition to a large number of immune-related processes, the positive regulation of fibroblast proliferation and the negative regulation of angiogenesis also appeared at M4. The processes related to phagocytosis and proteolysis began to appear at M5.

**Figure 5.**
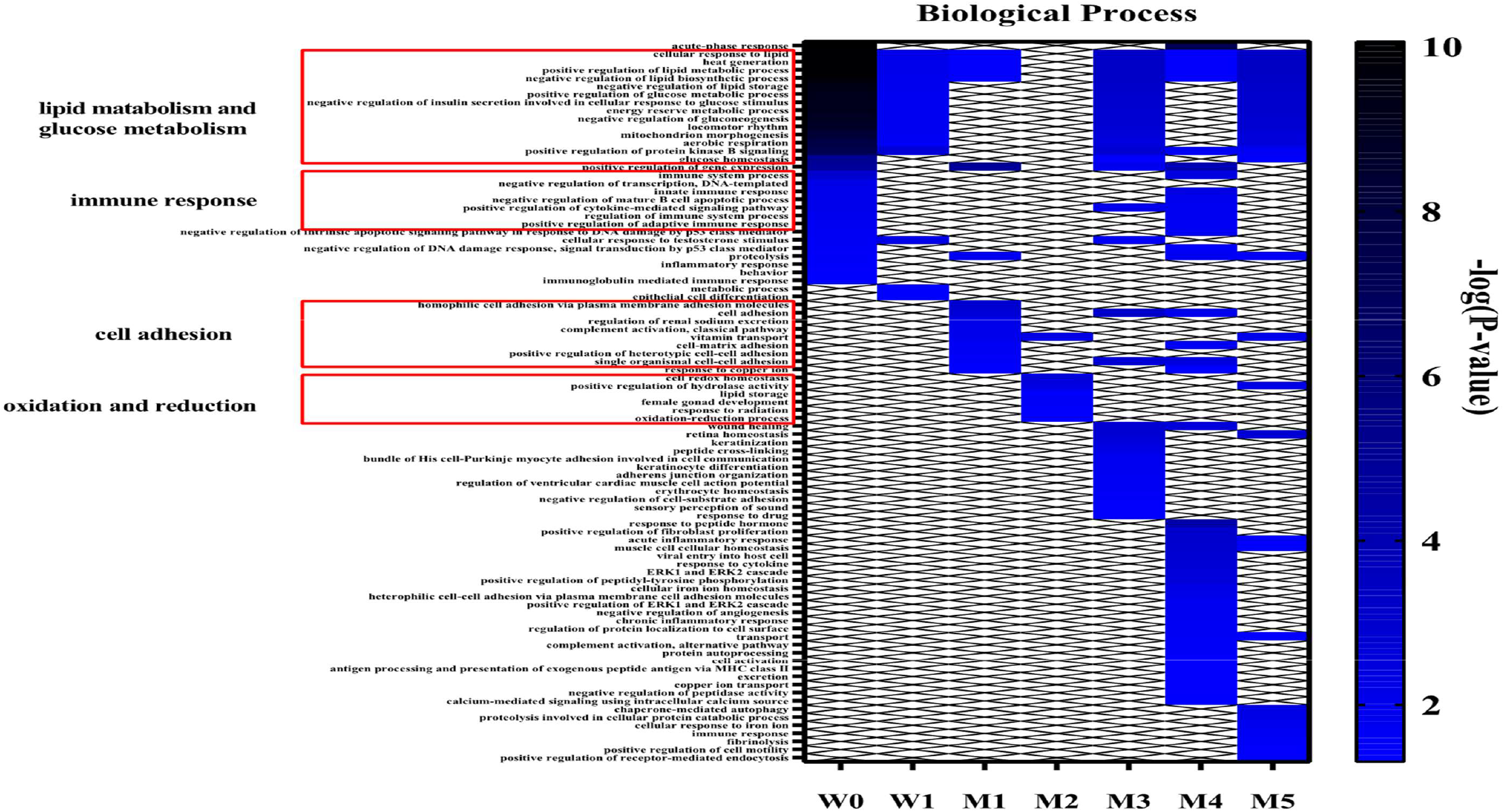
Functional annotation of differential proteins at different time points between the experimental and control groups (*p <*0.05)

##### (1) Effects of genetic factors

At W0, before a high-fat diet was administered to the experimental group, the only difference between the two groups was genetic factors. There were already significant differences in the biological processes related to lipid and glycometabolism, indicating that *ApoE* gene knockout greatly affects the lipid transport in mice in the experimental group, which is reflected by the urinary proteome very early. Acute phase reactions, immune responses, cytokines and proteolysis are also closely related to atherosclerosis ^[45-48]^.

##### (2) Urinary proteome changes during whole process

The literature shows that during the early stages of atherosclerosis, low-density lipoprotein (LDL) particles accumulate in the arterial intima, thereby being protected from plasma antioxidants, undergoing oxidation and other modifications and having proinflammatory and immunogenic properties. Classic monocytes circulating in the blood can exhibit anti-inflammatory functions and bind to the adhesion molecules expressed by activated endothelial cells to enter the inner membrane. Once in the inner membrane, monocytes can mature into macrophages, which express scavenger receptors that bind to lipoprotein particles and then become foam cells, finally forming the core of atherosclerotic plaques. T lymphocytes can also enter the inner membrane to regulate the functions of natural immune cells, endothelial cells and smooth muscle cells. The smooth muscle cells in the media can migrate to the inner membrane under the action of leukocytes to secrete extracellular matrix and form a fibrous cap ^[49]^. During the exploration of this experiment, at week 1, the differentially expressed proteins between the experimental and control groups were related to the differentiation of epithelial cells, and cell adhesion was enriched in M1 macrophages, which may be related to the adhesion of monocytes. Differential proteins between the experimental and control groups at M2 were related to biological processes related to redox, which may be related to the redox of LDL particles. Cell adhesion also changes at M3, which may involve the recruitment of phagocytes. Numerous immune-related biological processes changed in M4, indicating the participation of immune cells, such as T cells. The regulation of fibroblast proliferation may be related to the formation of fibrous caps. Enriched results revealed that proteolysis changed significantly at M5. It has been reported that activated macrophages can secrete proteolytic enzymes and degrade matrix components. The loss of matrix components may subsequently lead to plaque instability and increase the risk of plaque rupture and thrombosis ^[50]^. Fibrin dissolution also plays an important role in the development of atherosclerosis ^[51]^.

The biological processes of the enrichment of differential proteins at different time points can correspond to the occurrence and development of atherosclerosis, indicating that the urinary proteome has the potential to be used to monitor the disease process.

After a week of a high-fat diet in the experimental group, the protein kinase B signalling pathway changed. It is reported to play an important role in the survival, proliferation and migration of macrophages and may affect the development of atherosclerosis ^[52]^. After a month of a high-fat diet, many biological processes underwent significant changes. Studies have shown that urinary sodium excretion is the decisive factor of carotid intima-media thickness, which is an indicator of atherosclerosis ^[53]^. The classical pathway of complement activation is also related to atherosclerosis ^[54]^. Copper and isotype cysteine can interact to generate free radicals, thereby oxidizing LDL, which has been found in atherosclerotic plaques ^[55]^. At M2, oestrogen has been reported to have a variety of anti-atherosclerotic properties, including affecting plasma lipoprotein levels and stimulating the production of prostacyclin and nitric oxide ^[56]^. At M3, wound healing is also associated with atherosclerosis ^[40]^. For the biological processes changed at M4, the ERK1/ERK2 pathway plays an important role in the proliferation of vascular smooth muscle cells induced by insulin (INS) and thrombin ^[43]^. Alternative pathways of complement activation and major histocompatibility complex family II have been reported to be associated with atherosclerosis ^[57, 58]^. In the enrichment of differential proteins at M5, chaperone-mediated autophagy (CMA) plays an important upstream regulatory role in lipid metabolism and lipid metabolism in liver cells ^[59]^.

When the experimental group was compared to the control group, there was a large difference in W0, demonstrating that the urinary proteome reflects even slight difference between the groups. In the subsequent control results at each time point, the degree of overlap in the differential proteins was small, but they were mostly related to lipids and cardiovascular diseases. The enriched biological processes also correspond to the progression of atherosclerosis, indicating that the urinary proteome is useful to monitor the disease process. However, as mentioned before, this type of comparison does not take the influence of diet and other factors into account.

### 3.3 Chemical modifications of proteins

To further explore the effect of high-fat diet on chemical modifications of urine proteins, a total of 15 samples were selected at three time points (EW0/EM5/CM5). After data retrieval (.raw) based on open-pFind software, the analysis results were exported in pBuild.

A total of 923 different chemical modification types were identified in 15 samples, of which 468 chemical modification types were identified in the EW0 group, 748 chemical modification types were identified in the EM5 group, and 611 chemical modification types were identified in the CM5 group.

An unsupervised cluster analysis of all modifications found that the CM5 group was well distinguished from the other two groups (Figure 6). The percentages of different modification types in the EW0 group and the EM5 group were quantified to identify the modification changes that occurred in the internal controls. Among them, one modification type was unique to the EW0 group and existed in more than 4 samples (the total number of samples was 5), 23 modification types were unique to the EM5 group and existed in more than 5 samples (the total number of samples was 6), there are 68 types of modifications shared by the two groups, and there were significant differences (FC≥1.5 or ≤0.67, *p <*0.05). At the same time, the proportions of different types of modified sites in the CM5 group and the EM5 group were quantified, and the difference between the experimental group and the control group was analysed. Among them, 8 modification types were unique to the CM5 group and existed in more than 3 samples (the total number of samples was 4), and 19 modification types were unique to the EM5 group and existed in more than 5 samples (the total number of samples was 6). There were 72 types of modifications that were shared by the two groups, and there were significant differences (FC≥1.5 or ≤0.67, *p <*0.05) (see Table S5 for details).

**Figure 6.**
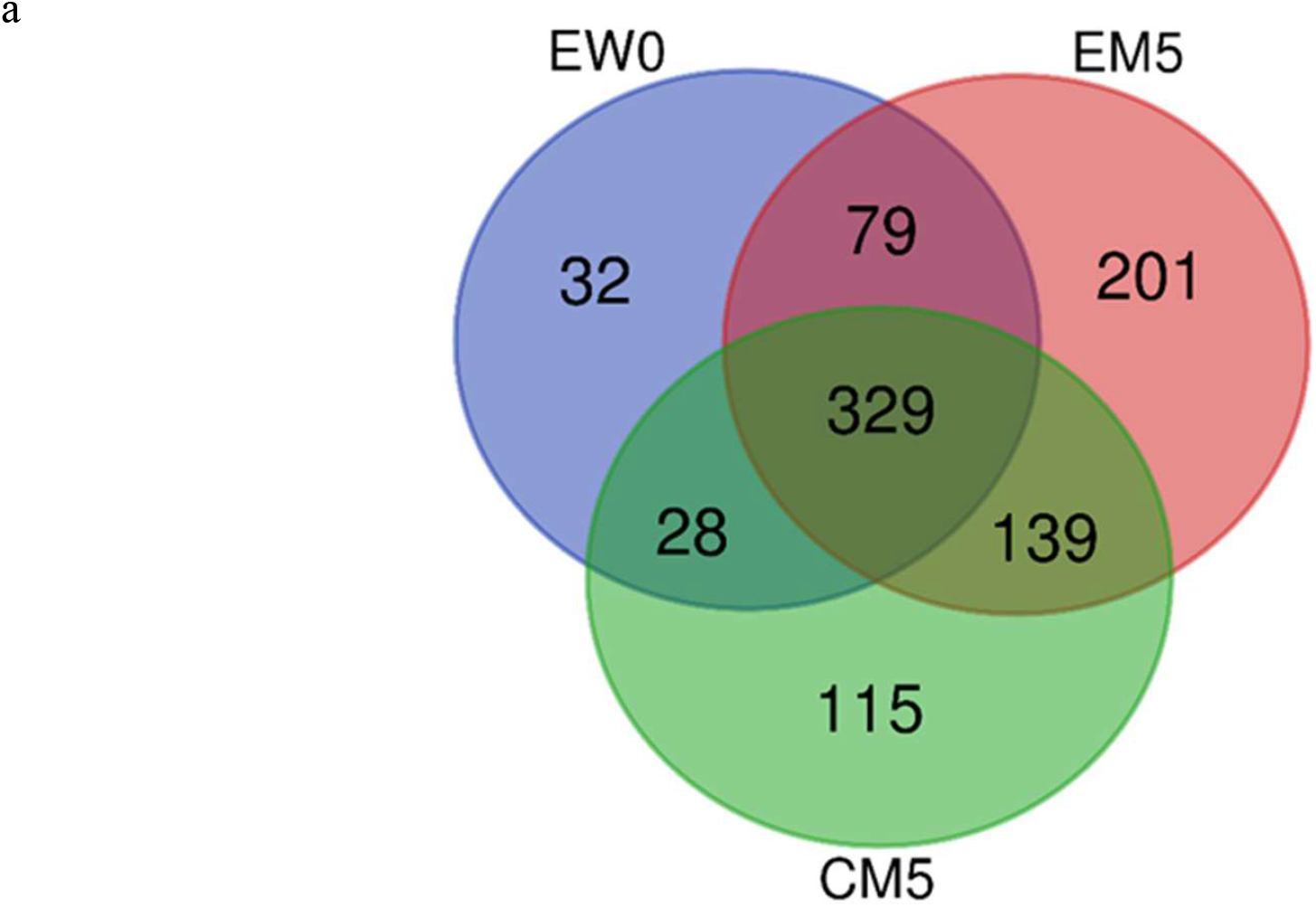
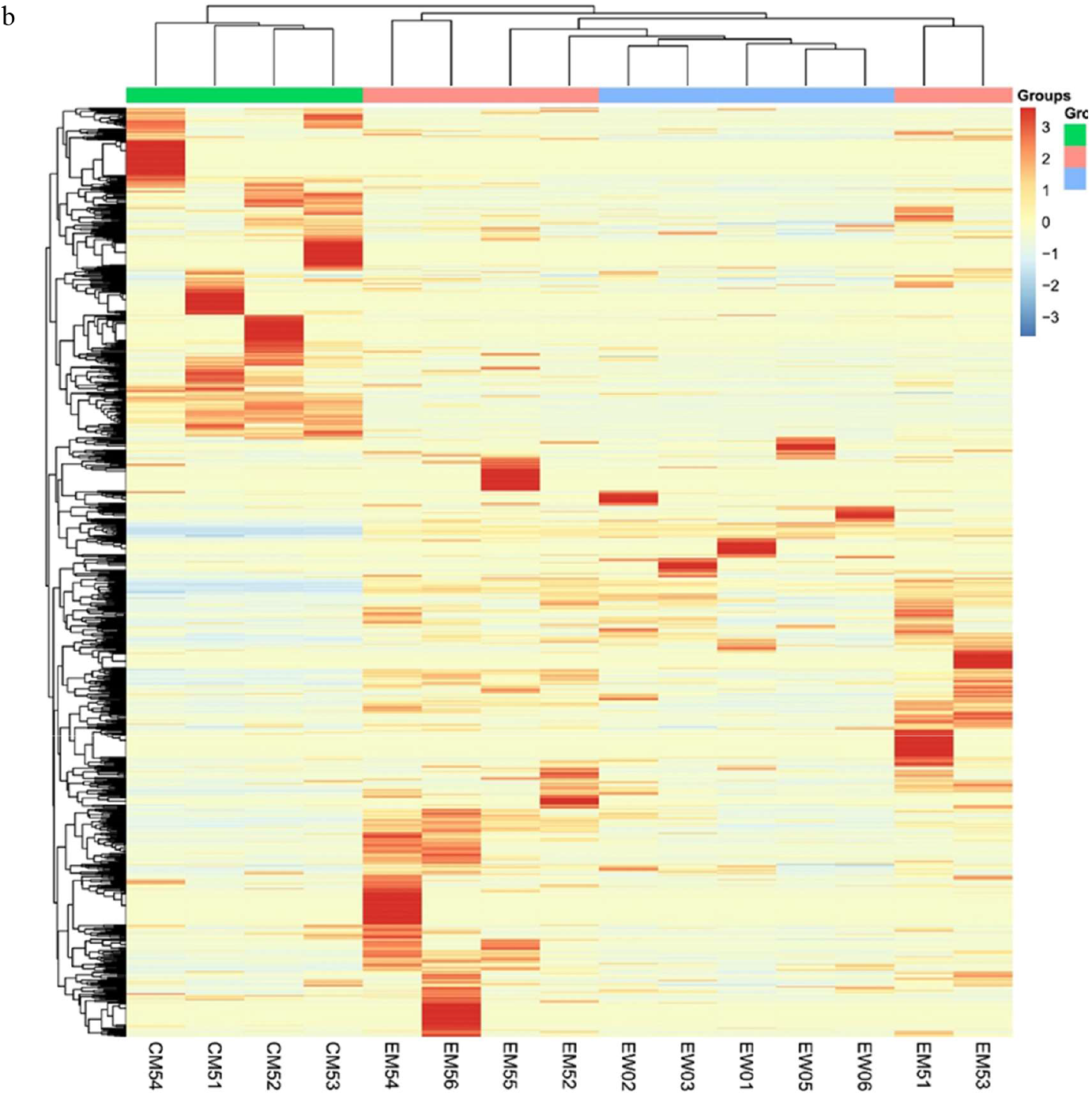
Chemical modifications among the three groups. (a) Venn diagram of modification types among the three groups. (b) Unsupervised clustering of all the modification types in the three groups.

To reduce the false negative influence caused by the open search mode, a restricted search method was used for verification. Modification types that accounted for the top five modification sites in the open search were fixed, modification types that were unique in a group and existed in each sample and modification types that have been reported related to lipids in the literature are selected. Twenty modifications in the EM5-EW0 group and 25 modifications in the EW5-CM5 group were selected, and the proportion of modified sites in the total number of sites was calculated (Table S6). The screening criteria were FC≥1.5 or ≤ 0.67 and *p <*0.05. Finally, in the internal control of the experimental group (EM5-EW0), the N-terminal carbamylation (Carbamyl[AnyN-term]), the CHDH modification of aspartic acid (CHDH[D]), the tryptophan by kynurenine acid substitution (Trp->Kynurenin[W]), oxidation modification of proline (Oxidation[P]), cysteine modification of cysteine (Cysteinyl[C]), sulfur dioxide modification of cysteine (SulfurDioxide[C]), the NO_SMX_SIMD modification of cysteine (NO_SMX_SIMD[C]) and the Delta_H(2)C(3) modification of lysine (Delta_H(2)C(3)[K]) significantly changed. In the comparison between the experimental and control groups (EM5-CM5), the guanidine modification of lysine (Guanidinyl[K]), the phosphouridine modification of tyrosine (PhosphoUridine[Y]), the N-terminal carbamoyl Modification (Carbamyl[AnyN-term]), Delta_H(2)C(2) modification (Delta_H(2)C(2)[AnyN-term]) at the N-terminus and Dihydroxyimidazolidine modification of arginine (Dihydroxyimidazolidine) [R]) showed significant changes.

Among the changes observed in the internal control of the experimental group, many studies have shown that carbamylated proteins are involved in the occurrence of diseases, especially atherosclerosis and chronic renal failure ^[136]^. The kynurenine pathway is the primary pathway of tryptophan metabolism and plays an important role in early atherosclerosis ^[137]^. The oxidation of proline can form glutamate semialdehyde, and glutamate semialdehyde is closely related to lipid peroxidation^[138]^. Elevated plasma homocysteine has also been widely studied as an independent risk factor for atherosclerosis ^[139]^. Obstruction of the sulfur dioxide/aspartate aminotransferase pathway is also known to be involved in the pathogenesis of many cardiovascular diseases ^[140]^. The Delta_H(2)C(3) modification of lysine also refers to acrolein addition +38, and acrolein and other α- and β-unsaturated aldehydes are considered to be mediators of inflammation and vascular dysfunction ^[141]^. CHDH modification of aspartic acid and NO_SMX_SIMD modification of cysteine have not been reported to be related to atherosclerosis and may act as potential modification sites.

Although it was not verified in a restricted search, there are also studies claiming that the interruption of cell signals mediated by electrophiles is related to the occurrence of atherosclerosis and cancer. HNE and ONE and their derivatives are both active lipid electrophile reagents that inhibit the release of proinflammatory factors to a certain extent ^[142]^. Nε-carboxymethyl-lysine (CML) has been reported to accumulate in large amounts in the tissues of diabetes and atherosclerosis, and glucosone aldehyde is related to its formation ^[143]^. Benzyl isothiocyanate salt has been reported to inhibit lipid production and fatty liver formation in obese mice fed a high-fat diet ^[144]^. It has been reported in the literature that thiazolidine derivatives have a positive effect in the treatment of LDLR(-/-) atherosclerotic mice ^[145]^. In addition, the carboxyethylation of lysine has also changed, and some literature indicates that the degree of carboxymethylation and carboxyethylation of lysine in the plasma of diabetic mice is significantly increased ^[146]^. Changes in the expression of fucosylated oligosaccharides have been observed in pathological processes such as atherosclerosis ^[147]^. In addition, the phosphorylation modification of tyrosine is related to the formation of esters, which may also be involved in lipid metabolism and the occurrence and progression of diseases.

In the differential modifications between the experimental group and the control group, some of the significantly changed modifications also changed in the internal control of the experimental group. In addition, the Delta_H(2)C(2) modification at the N-terminus of the amino acid also refers to acetaldehyde +26. In addition, acetaldehyde stimulates the growth of vascular smooth muscle cells in a notch-dependent manner, promoting the occurrence of atherosclerosis ^[148]^. Advanced protein glycosylation is an important mechanism for the development of advanced complications of diabetes, including atherosclerosis. Hydroimidazolone-1 derived from methylglyoxal is the most abundant advanced glycosylation end product in human plasma ^[149]^. In addition, the guanidine modification of lysine may also be related to atherosclerosis.

Although did not been verified in the restricted search, an increasing number of studies have shown that short-chain fatty acids and their homologous acylation are involved in cardiovascular disease, and the proportions of 2-hydroxyisobutyrylation, malonylation and crotonylation in the experimental group were significantly increased^[150]^. Nε-carboxymethyl-lysine (CML) has been reported to accumulate in large amounts in the tissues in diabetes and atherosclerosis, and its induced PI3K/Akt signal inhibition promotes foam cell apoptosis and the progression of atherosclerosis ^[151]^. In addition, glucosone is closely related to its formation, the proportion of which also increased significantly in the experimental group. Oxidation of tyrosine produces dihydroxyphenylalanine (DOPA), and the protein binding DOPA in tissues is elevated in many age-related pathological diseases, such as atherosclerosis and cataract formation ^[152]^.

As mentioned above, the internal controls of the experimental group avoid the influence of genes, diet and other factors on the urinary proteome, but they may be affected by the growth and development of the organisms themselves. Intergroup control avoids the influence of development but cannot avoid factors such as diet. The identification results of chemical modifications of urine proteins showed that regardless of whether internal control or intergroup control was adopted, the modification status changed greatly and was closely related to lipids and cardiovascular diseases. In comparison, differences in the intergroup control may be more obvious.

## 4 Conclusion

This study explored changes in urinary proteomics of high-fat diet-fed *ApoE*^-/-^ mice. The internal control results showed that even after only one week of a high-fat diet, when the urinary proteome of the control group was not significantly changed, the urinary proteome of the experimental group had changed significantly, and most of the enriched biological pathways were related to lipid metabolism and glycometabolism, indicating that the urinary proteome has the potential for early and sensitive monitoring of biological changes. Most of the proteins and their family members that change continually in disease progression have been reported to be related to cardiovascular diseases or can be used as biomarkers. The results of the intergroup comparison showed that the biological processes enriched by differential proteins at different time points correspond to the occurrence and development of the atherosclerosis, indicating that the urinary proteome has the potential to be used to monitor the disease process. The differential modification types in the internal control of the experimental group and the comparison between the experimental and control groups have also been reported to be related to lipids and cardiovascular diseases, which can be used as a reference for identifying new biomarkers.

## Declaration

### Ethics approval and consent to participate

All animal protocols governing the experiments in this study were approved by the Institute of Basic Medical Sciences Animal Ethics Committee, Peking Union Medical College (Approved ID: ACUC-A02-2014-007).

### Consent for publication

Not applicable.

### Availability of data and material

The mass spectrometry proteomics data have been deposited to the ProteomeXchange Consortium (http://proteomecentral.proteomexchange.org) via the iProX partner repository with the dataset identifier PXD027610.

### Authors’ contributions

**Hua Yuanrui** performed the experiments, analyzed the data, contributed reagents/materials/ analysis tools, prepared figures and/or tables, authored or reviewed drafts of the paper, approved the final draft. **Meng Wenshu** and **Wei Jing** performed the experiments and contributed reagents/materials/analysis tools. **Liu Yongtao** analyzed the data and contributed reagents/materials/analysis tools. **Gao Youhe** conceived and designed the experiments, authored or reviewed drafts of the paper, approved the final draft.

## Acknowledgements

Not applicable.

