## Supplementary material for "Changes of urinary proteome in high-fat diet *ApoE*^-/-^ mice": Table S1

| Group | Total number of random combinations | Average numbers of proteins with false random combinations | Number of correctly identified differential proteins | Percentage |
|---|---|---|---|---|
| EW1-EW0 | 231 | 8.47 | 27 | 31.37% |
| EM1-EW0 | 231 | 10.35 | 51 | 20.29% |
| EM2-EW0 | 231 | 9.99 | 69 | 14.48% |
| EM3-EW0 | 231 | 9.69 | 86 | 11.27% |
| EM4-EW0 | 231 | 8.67 | 65 | 13.33% |
| EM5-EW0 | 231 | 9.64 | 88 | 10.95% |
| EW0-CW0 | 126 | 9.66 | 54 | 17.89% |
| EW1-CW1 | 210 | 9.07 | 17 | 53.35% |
| EM1-CM1 | 210 | 8.31 | 55 | 15.11% |
| EM2-CM2 | 210 | 10.16 | 25 | 40.64% |
| EM3-CM3 | 210 | 9.24 | 58 | 15.93% |
| EM4-CM4 | 210 | 9.48 | 62 | 15.29% |
| EM5-CM5 | 210 | 8.53 | 55 | 15.51% |

Table S1 Screening results of random combinations of urine samples

To test the influence of the possibility of random generation of different proteins on the screening results, every two groups of samples were randomly arranged and combined, and the average number of different proteins in all combinations was calculated according to the same screening conditions (Fc ≥1.5 or Fc ≤0.67, $p$ <0.05). The average number of differential proteins produced by random grouping was significantly different from the number of differential proteins obtained by normal grouping screening, indicating that the differential proteins were indeed generated by the differences in samples within the group, which was relatively reliable.

| UniProt ID | Protein Name | P-value | Fold Change |
|---|---|---|---|
| O35887 | Calumenin | 0.0386 | 2.52 |
| Q60932 | Voltage-dependent anion-selective channel protein 1 | 0.0382 | 1.82 |
| P01660 | Ig kappa chain V-III region PC 3741/TEPC 111 | 0.0458 | 1.64 |
| Q8K426 | Resistin-like gamma | 0.0119 | 1.59 |
| P01898 | H-2 class I histocompatibility antigen, Q10 alpha chain | 0.0290 | 0.67 |
| P23953 | Carboxylesterase 1C | 0.0333 | 0.61 |
| Q60847 | Collagen alpha-1(XII) chain | 0.0437 | 0.56 |
| Q07456 | Protein AMBP | 0.0068 | 0.51 |
| O88322 | Nidogen-2 | 0.0063 | 0.51 |
| Q9JJS0 | Signal peptide, CUB and EGF-like domain-containing protein 2 | 0.0135 | 0.46 |
| P09036 | Serine protease inhibitor Kazal-type 1 | 0.0443 | 0.44 |

Table S2 Differential proteins between week 1 and week 0 samples in the control group.

| Human UniProt | Protein Name | Fold Change | | | | | | R |
|---|---|---|---|---|---|---|---|---|
| | | EW0 | EM1 | EM2 | EM3 | EM4 | EM5 | |
| No | Major urinary protein 17 | 1 | 11.94 | 27.92 | 27.91 | 20.90 | 11.18 | |
| No | Major urinary protein 1 | 1 | 9.59 | 18.11 | 17.19 | 21.96 | 9.24 | |
| No | Ig kappa chain V-III region PC 7043 | 1 | — | 11.01 | 8.41 | 18.15 | 9.95 | |
| P00738 | Haptoglobin | 1 | — | 7.68 | 8.66 | 5.76 | 11.25 | |
| P04233 | H-2 class II histocompatibility antigen gamma chain | 1 | — | 4.99 | 5.50 | 10.95 | 4.80 | |
| No | Major urinary protein 20 | 1 | 4.24 | 6.10 | 5.57 | 7.53 | 4.28 | |
| No | Major urinary protein 5 | 1 | 3.94 | 5.26 | 4.40 | 6.76 | 3.95 | |
| No | Lymphocyte antigen 6C2 | 1 | 2.91 | 4.45 | 2.95 | 6.45 | 2.59 | |
| Q03403 | Trefoil factor 2 | 1 | 1.76 | 3.04 | 2.16 | 5.91 | 2.41 | |
| O95998 | Interleukin-18-binding protein | 1 | — | 3.01 | 2.08 | 3.93 | 3.08 | |
| No | Major urinary protein 2 | 1 | — | 2.52 | 2.56 | 4.30 | 2.43 | |
| P13473 | Lysosome-associated membrane glycoprotein 2 | 1 | — | 2.10 | 3.13 | 3.20 | 2.64 | |
| O95497 | Pantetheinase | 1 | — | 1.73 | 1.79 | 1.75 | 5.22 | |
| P12830 | Cadherin-1 | 1 | — | 2.20 | 2.88 | 2.28 | 2.13 | |
| P61109 | Kidney androgen-regulated protein | 1 | — | 1.84 | 1.92 | 3.54 | 1.63 | |
| P01834 | Immunoglobulin kappa constant | 1 | — | 1.94 | 2.78 | 2.35 | 1.61 | |
| No | Ig kappa chain V-II region 26-10 | 1 | — | 1.74 | 1.78 | 2.77 | 1.85 | |
| Q01459 | Di-N-acetylchitobiase | 1 | — | 1.65 | 2.16 | 1.99 | 2.08 | |
| P02751 | Fibronectin | 1 | — | 0.63 | 0.48 | 0.59 | 0.40 | |
| P02766 | Transthyretin | 1 | — | 0.51 | 0.53 | 0.56 | 0.49 | |
| P02787 | Serotransferrin | 1 | — | 0.46 | 0.45 | 0.37 | 0.45 | |
| P01019 | Angiotensinogen | 1 | 0.44 | 0.47 | 0.46 | 0.37 | 0.35 | |
| P01011 | Serine protease inhibitor A3K | 1 | — | 0.38 | 0.38 | 0.35 | 0.37 | |
| P01009 | Alpha-1-antitrypsin 1-1 | 1 | 0.51 | 0.29 | 0.29 | 0.30 | 0.41 | |
| O95388 | CCN family member 4 | 1 | 0.24 | 0.30 | 0.40 | 0.42 | 0.20 | |
| P20062 | Transcobalamin-2 | 1 | 0.36 | 0.37 | 0.27 | 0.31 | 0.19 | |
| Q9P1F3 | Costars family protein ABRACL | 1 | 0.32 | 0.29 | 0.37 | 0.26 | 0.25 | |
| Q06141 | Regenerating islet-derived protein 3-beta | 1 | 0.18 | 0.27 | 0.25 | 0.37 | 0.32 | |
| P12273 | Prolactin-inducible protein homolog | 1 | — | 0.30 | 0.24 | 0.21 | 0.35 | |
| P02765 | Alpha-2-HS-glycoprotein | 1 | 0.39 | 0.27 | 0.16 | 0.32 | 0.11 | |
| P06396 | Gelsolin | 1 | 0.39 | 0.27 | 0.18 | 0.24 | 0.15 | |
| Q9NQ36 | Signal peptide, CUB and EGF-like domain-containing protein 2 | 1 | 0.23 | 0.24 | 0.17 | 0.23 | 0.09 | |
| Q14112 | Nidogen-2 | 1 | 0.24 | 0.20 | 0.14 | 0.29 | 0.08 | |
| Q6P5S2 | Protein LEG1 homolog | 1 | — | 0.16 | 0.11 | 0.08 | 0.11 | |
| P02452 | Collagen alpha-1(I) chain | 1 | 0.07 | 0.11 | 0.10 | 0.14 | 0.06 | |

Table S3 Details of continuously changing differential proteins in the internal control of the experimental group.

| Group | UniProt | Human UniProt | Protein Name | P-value | Fold Change |
|---|---|---|---|---|---|
| | P10287 | P22223 | Cadherin-3 | 0.0245 | 2.74 |
| | Q07797 | Q08380 | Galectin-3-binding protein | 0.0179 | 2.57 |
| | P23780 | P16278 | Beta-galactosidase | 0.0468 | 2.05 |
| | Q8K1H9 | Q9NY56 | Odorant-binding protein 2a | 0.0328 | 2.04 |
| | Q9WVJ3 | Q9Y646 | Carboxypeptidase Q | 0.0235 | 1.88 |
| EM2 | Q91X17 | P07911 | Uromodulin | 0.0074 | 1.86 |
| - | P55288 | P55287 | Cadherin-11 | 0.0395 | 1.80 |
| EM1 | Q03404 | Q03403 | Trefoil factor 2 | 0.0131 | 1.73 |
| | Q91WR6 | Q9NU53 | Glycoprotein integral membrane protein 1 | 0.0126 | 1.63 |
| | Q9ESY9 | P13284 | Gamma-interferon-inducible lysosomal thiol reductase | 0.0112 | 1.58 |
| | P06909 | P08603 | Complement factor H | 0.0192 | 0.61 |
| | Q8BND5 | O00391 | Sulfhydryl oxidase 1 | 0.0176 | 0.55 |
| | Q8K0E8 | P02675 | Fibrinogen beta chain | 0.0329 | 0.24 |
| | Q00993 | P30530 | Tyrosine-protein kinase receptor UFO | 0.0114 | 2.05 |
| | Q62267 | No | Cornifin-B | 0.0153 | 1.89 |
| | P70663 | Q14515 | SPARC-like protein 1 | 0.0128 | 1.79 |
| EM3 | P70699 | P10253 | Lysosomal alpha-glucosidase | 0.0364 | 1.68 |
| - | P03953 | P00746 | Complement factor D | 0.0176 | 0.64 |
| EM2 | Q00898 | P01009 | Alpha-1-antitrypsin 1-5 | 0.0487 | 0.64 |
| | Q9Z0L8 | Q92820 | Gamma-glutamyl hydrolase | 0.0456 | 0.54 |
| | Q60847 | Q99715 | Collagen alpha-1(XII) chain | 0.0499 | 0.51 |
| | Q9EP95 | Q9BQ08 | Resistin-like alpha | 0.0201 | 0.26 |
| | P25119 | P20333 | Tumour necrosis factor receptor superfamily member 1B | 0.0491 | 3.19 |
| | Q03404 | Q03403 | Trefoil factor 2 | 0.0237 | 2.74 |
| | Q9DAU7 | Q14508 | WAP four-disulfide core domain protein 2 | 0.0293 | 2.61 |
| | Q9EQC7 | O95633 | Follistatin-related protein 3 | 0.0460 | 2.58 |
| | Q07797 | Q08380 | Galectin-3-binding protein | 0.0410 | 2.57 |
| | Q61581 | Q16270 | Insulin-like growth factor-binding protein 7 | 0.0404 | 2.50 |
| EM4 | Q01339 | P02749 | Beta-2-glycoprotein 1 | 0.0411 | 2.45 |
| - | Q6SJQ5 | Q6UXZ3 | CMRF35-like molecule 3 | 0.0430 | 2.28 |
| EM3 | P15379 | P16070 | CD44 antigen | 0.0313 | 2.18 |
| | Q91WR8 | P59796 | Glutathione peroxidase 6 | 0.0096 | 1.99 |
| | Q9Z0M9 | O95998 | Interleukin-18-binding protein | 0.0294 | 1.89 |
| | Q5SSE9 | Q86UQ4 | ATP-binding cassette sub-family A member 13 | 0.0279 | 1.69 |
| | P01132 | P01133 | Pro-epidermal growth factor | 0.0125 | 1.69 |
| | P01631 | No | Ig kappa chain V-II region 26-10 | 0.0087 | 1.56 |
| | P07724 | P02768 | Albumin | 0.0193 | 0.57 |
| | P01864 | No | Ig gamma-2A chain C region secreted form | 0.0121 | 3.35 |
| EM5 | Q9ET22 | Q9UHL4 | Dipeptidyl peptidase 2 | 0.0002 | 3.31 |
| - | Q9Z0K8 | O95497 | Pantetheinase | 0.0160 | 2.98 |
| EM4 | P00688 | P04746 | Pancreatic alpha-amylase | 0.0072 | 2.76 |
| | P13634 | P00915 | Carbonic anhydrase 1 | 0.0017 | 2.51 |

| | | | | |
|---|---|---|---|---|
| A2ARV4 | P98164 | Low-density lipoprotein receptor-related protein 2 | 0.0248 | 2.11 |
| P51910 | P05090 | Apolipoprotein D | 0.0496 | 2.04 |
| Q9WUU7 | Q9UBR2 | Cathepsin Z | 0.0121 | 1.78 |
| P16675 | P10619 | Lysosomal protective protein | 0.0190 | 1.77 |
| P07724 | P02768 | Albumin | 0.0094 | 1.68 |
| Q06890 | P10909 | Clusterin | 0.0384 | 1.52 |
| Q61147 | P00450 | Ceruloplasmin | 0.0491 | 1.52 |
| P01631 | No | Ig kappa chain V-II region 26-10 | 0.0067 | 0.67 |
| P61110 | P61109 | Kidney androgen-regulated protein | 0.0270 | 0.46 |
| Q61129 | P05156 | Complement factor I | 0.0213 | 0.44 |
| P04441 | P04233 | H-2 class II histocompatibility antigen gamma chain | 0.0483 | 0.44 |
| Q01339 | P02749 | Beta-2-glycoprotein 1 | 0.0439 | 0.42 |
| P42567 | P42566 | Epidermal growth factor receptor substrate 15 | 0.0482 | 0.41 |
| Q03404 | Q03403 | Trefoil factor 2 | 0.0317 | 0.41 |
| Q91WR8 | P59796 | Glutathione peroxidase 6 | 0.0021 | 0.38 |
| Q61271 | P36896 | Activin receptor type-1B | 0.0275 | 0.35 |
| Q62395 | Q07654 | Trefoil factor 3 | 0.0402 | 0.34 |
| P29699 | P02765 | Alpha-2-HS-glycoprotein | 0.0484 | 0.34 |
| Q08423 | P04155 | Trefoil factor 1 | 0.0447 | 0.32 |
| P10287 | P22223 | Cadherin-3 | 0.0126 | 0.31 |
| Q9EP95 | Q9BQ08 | Resistin-like alpha | 0.0479 | 0.30 |
| O08997 | O00244 | Copper transport protein ATOX1 | 0.0378 | 0.28 |
| P0CG49 | P0CG47 | Polyubiquitin-B | 0.0095 | 0.28 |
| O88322 | Q14112 | Nidogen-2 | 0.0196 | 0.28 |
| Q7TMJ8 | Q96FE7 | Phosphoinositide-3-kinase-interacting protein 1 | 0.0360 | 0.26 |
| O09051 | Q16661 | Guanylate cyclase activator 2B | 0.0252 | 0.23 |
| P09036 | P00995 | Serine protease inhibitor Kazal-type 1 | 0.0200 | 0.19 |
| Q62267 | No | Cornifin-B | 0.0500 | 0.19 |

Table S4 Differential proteins between adjacent time points of the experimental group.

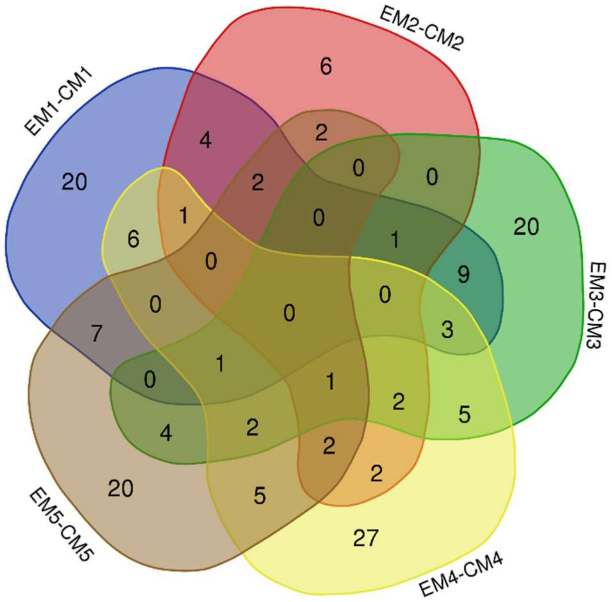

Figure S1 Venn diagram of differential proteins at different time points between the experimental group and the control group.

| Group | Modification Name | Modification Type | P-value | Fold Change |
|---|---|---|---|---|
| | Oxidation[P] | Post-translational | ① | ① |
| | 4-ONE[H] | Chemical derivative | ② | ② |
| | AccQTag[AnyN-term] | Chemical derivative | ② | ② |
| | AEBS[K] | Artefact | ② | ② |
| | Cation_K[D] | Artefact | ② | ② |
| | Delta_H(2)C(3)[K] | Other | ② | ② |
| | Ethyl+Deamidated[N] | Chemical derivative | ② | ② |
| | Hep[T] | O-linked glycosylation | ② | ② |
| | NHS-LC-Biotin[AnyN-term] | Chemical derivative | ② | ② |
| | NO_SMX_SIMD[C] | Chemical derivative | ② | ② |
| | Arg->Ser[R] | AA substitution | ② | ② |
| | CHDH[D] | Post-translational | ② | ② |
| | Diisopropylphosphate[AnyN-term] | Chemical derivative | ② | ② |
| | glucosone[R] | Other | ② | ② |
| | Gly->His[G] | AA substitution | ② | ② |
| | Hex[K] | Other glycosylation | ② | ② |
| | ICDID[C] | Artefact | ② | ② |
| | ICPL_2H(4)[AnyN-term] | Artefact | ② | ② |
| EM5 | LG-anhydrolactam[AnyN-term] | Post-translational | ② | ② |
| - | NDA[K] | Chemical derivative | ② | ② |
| EW0 | Phe->Gly[F] | AA substitution | ② | ② |
| | Phospho[C] | Post-translational | ② | ② |
| | Phosphoadenosine[T] | Post-translational | ② | ② |
| | SMA[AnyN-term] | Chemical derivative | ② | ② |
| | BITC[AnyN-term] | Chemical derivative | ② | ② |
| | Bromo[F] | Post-translational | ② | ② |
| | Cytopiloyne+water[AnyN-term] | Chemical derivative | ② | ② |
| | DAET[S] | Chemical derivative | ② | ② |
| | Delta_Hg(1)[C] | Chemical derivative | ② | ② |
| | Didehydro[Y] | Post-translational | ② | ② |
| | Dimethylphosphothione[C] | Chemical derivative | ② | ② |
| | Glu->Tyr[E] | AA substitution | ② | ② |
| | Hex(2)[T] | O-linked glycosylation | ② | ② |
| | HexNAc(2)Sulf(1)[S] | O-linked glycosylation | ② | ② |
| | HNE+Delta_H(2)[K] | Chemical derivative | ② | ② |
| | Iminobiotin[AnyN-term] | Chemical derivative | ② | ② |
| | Iodoacetanilide_13C(6)[AnyN-term] | Artefact | ② | ② |
| | Nitro[F] | Artefact | ② | ② |
| | Oxidation[V] | Chemical derivative | ② | ② |
| | Oxidation+NEM[C] | Chemical derivative | ② | ② |

| | | | |
|---|---|---|---|
| PhosphoUridine[Y] | Post-translational | ② | ② |
| Thiophospho[T] | Other | ② | ② |
| Xle->Phe[I] | AA substitution | ② | ② |
| Xlink_SMCC[219][C] | Chemical derivative | ② | ② |
| Xlink_SMCC[237][C] | Chemical derivative | ② | ② |
| NEIAA[C] | Artefact | 28.64 | 0.0378 |
| ICPL_13C(6)[AnyN-term] | Artefact | 21.41 | 0.0151 |
| Xle->Trp[I] | AA substitution | 14.55 | 0.0002 |
| Thiazolidine[W] | Chemical derivative | 14.17 | 0.0194 |
| SMA[K] | Chemical derivative | 13.45 | 0.0082 |
| NEMsulfurWater[C] | Chemical derivative | 13.24 | 0.0000 |
| Guanidinyl[K] | Chemical derivative | 13.08 | 0.0009 |
| Phosphopropargyl[S] | Multiple | 12.72 | 0.0348 |
| Thiophospho[S] | Other | 12.15 | 0.0007 |
| phenylsulfonylethyl[C] | Chemical derivative | 11.97 | 0.0013 |
| Ub-Br2[C] | Chemical derivative | 10.05 | 0.0106 |
| Asn->Trp[N] | AA substitution | 9.40 | 0.0035 |
| NEMsulfur[C] | Chemical derivative | 8.81 | 0.0002 |
| Carboxyethyl[K] | Post-translational | 7.43 | 0.0086 |
| Ethanedithiol[T] | Chemical derivative | 7.42 | 0.0247 |
| Met->Glu[M] | AA substitution | 7.11 | 0.0250 |
| DeStreak[C] | Chemical derivative | 6.62 | 0.0200 |
| Xlink_EGS[115][K] | Chemical derivative | 6.39 | 0.0015 |
| Xle->Tyr[L] | AA substitution | 6.26 | 0.0012 |
| 2-succinyl[C] | Chemical derivative | 6.07 | 0.0070 |
| Cys->SecNEM_2H(5)[C] | Chemical derivative | 5.80 | 0.0025 |
| Phenylisocyanate_2H(5)[AnyN-term] | Chemical derivative | 5.18 | 0.0054 |
| Asn->His[N] | AA substitution | 5.13 | 0.0316 |
| SulfurDioxide[C] | Post-translational | 4.64 | 0.0410 |
| Piperidine[AnyN-term] | Chemical derivative | 4.58 | 0.0379 |
| Tris[N] | Artefact | 4.47 | 0.0046 |
| Delta_H(-4)O(2)[W] | Chemical derivative | 4.42 | 0.0341 |
| Asn->Cys[N] | AA substitution | 4.35 | 0.0222 |
| Carboxymethyl[AnyN-term] | Artefact | 4.13 | 0.0371 |
| Xle->Tyr[I] | AA substitution | 4.07 | 0.0054 |
| Propionamide[C] | Artefact | 3.97 | 0.0065 |
| Xle->Cys[I] | AA substitution | 3.88 | 0.0410 |
| Carboxymethyl[K] | Artefact | 3.72 | 0.0001 |
| Isopropylphospho[T] | Chemical derivative | 3.72 | 0.0039 |
| Cation_Ni[II][D] | Artefact | 3.63 | 0.0468 |
| Trp->Kynurenin[W] | Chemical derivative | 3.56 | 0.0113 |
| 2-monomethylsuccinyl[C] | Chemical derivative | 3.55 | 0.0012 |
| PET[T] | Chemical derivative | 3.46 | 0.0499 |

| | | | | |
|---|---|---|---|---|
| | Cysteinyl[C] | Multiple | 3.44 | 0.0059 |
| | Nmethylmaleimide+water[C] | Chemical derivative | 3.43 | 0.0001 |
| | MDCC[C] | Chemical derivative | 3.40 | 0.0094 |
| | MercaptoEthanol[T] | Chemical derivative | 3.34 | 0.0196 |
| | Acetyl[AnyN-term] | Multiple | 3.28 | 0.0016 |
| | Formylasparagine[H] | Chemical derivative | 3.26 | 0.0100 |
| | Glucuronyl[T] | O-linked glycosylation | 3.10 | 0.0481 |
| | O-Methylphosphate[S] | Chemical derivative | 2.87 | 0.0318 |
| | Trioxidation[C] | Chemical derivative | 2.86 | 0.0036 |
| | Propionamide[AnyN-term] | Chemical derivative | 2.85 | 0.0399 |
| | Asn->Met[N] | AA substitution | 2.81 | 0.0195 |
| | Carbamyl[AnyN-term] | Multiple | 2.75 | 0.0008 |
| | Cation_Ni[II][E] | Artefact | 2.65 | 0.0437 |
| | Cation_Fe[II][E] | Artefact | 2.64 | 0.0114 |
| | Trp->Oxolactone[W] | Chemical derivative | 2.49 | 0.0071 |
| | Arg[AnyN-term] | Other | 2.44 | 0.0268 |
| | dHex[N] | N-linked glycosylation | 2.32 | 0.0048 |
| | Amidine[AnyN-term] | Chemical derivative | 2.24 | 0.0444 |
| | Carboxy[W] | Chemical derivative | 2.14 | 0.0283 |
| | Nmethylmaleimide[C] | Chemical derivative | 2.06 | 0.0037 |
| | Dethiomethyl[M] | Artefact | 1.96 | 0.0295 |
| | Unknown_210[AnyN-term] | Artefact | 1.84 | 0.0000 |
| | Gly->Pro[G] | AA substitution | 1.76 | 0.0143 |
| | Formyl[T](Thr->Glu[T]) | Artefact | 1.68 | 0.0161 |
| | Xle->Gln[I] | AA substitution | 1.55 | 0.0318 |
| | CarbamidomethylDTT[C] | Artefact | 0.45 | 0.0008 |
| | Pyro-carbamidomethyl[AnyN-termC] | Artefact | 0.39 | 0.0011 |
| | Thr->Ala[T] | AA substitution | 0.39 | 0.0087 |
| | Deoxy[S](Ser->Ala[S]) | Chemical derivative | 0.36 | 0.0006 |
| | Gln->pyro-Glu[AnyN-termQ] | Artefact | 0.34 | 0.0000 |
| | Arg->Trp[R] | AA substitution | ③ | ③ |
| | His->Ser[H] | AA substitution | ③ | ③ |
| | HexNAc(2)[N] | N-linked glycosylation | ③ | ③ |
| | HN2_mustard[C] | Post-translational | ③ | ③ |
| | Oxidation[P] | Post-translational | ③ | ③ |
| EM5 - CM5 | Pro->His[P] | AA substitution | ③ | ③ |
| | Val->Glu[V] | AA substitution | ③ | ③ |
| | Xle->Val[I] | AA substitution | ③ | ③ |
| | Delta_H(2)C(3)[K] | Other | ② | ② |
| | DeStreak[C] | Chemical derivative | ② | ② |
| | Dihydroxyimidazolidine[R] | Multiple | ② | ② |
| | Guanidinyl[K] | Chemical derivative | ② | ② |

| | | | |
|---|---|---|---|
| phenylsulfonylethyl[C] | Chemical derivative | ② | ② |
| Cation_Cu[I][E] | Artefact | ② | ② |
| glucosone[R] | Other | ② | ② |
| Hex[K] | Other glycosylation | ② | ② |
| HexNAc(1)dHex(1)[T] | O-linked glycosylation | ② | ② |
| ICDID[C] | Artefact | ② | ② |
| ICPL_2H(4)[AnyN-term] | Artefact | ② | ② |
| LG-anhydrolactam[AnyN-term] | Post-translational | ② | ② |
| mTRAQ[AnyN-term] | Artefact | ② | ② |
| NDA[K] | Chemical derivative | ② | ② |
| Phe->Gly[F] | AA substitution | ② | ② |
| Quinone[W] | Post-translational | ② | ② |
| Quinone[Y] | Post-translational | ② | ② |
| SMA[K] | Chemical derivative | ② | ② |
| Thiazolidine[F] | Chemical derivative | ② | ② |
| 2-hydroxyisobutyrylation[K] | Post-translational | ② | ② |
| BITC[K] | Chemical derivative | ② | ② |
| Bromo[F] | Post-translational | ② | ② |
| Cytopiloyne+water[AnyN-term] | Chemical derivative | ② | ② |
| DAET[S] | Chemical derivative | ② | ② |
| Delta_Hg(1)[C] | Chemical derivative | ② | ② |
| HexNAc(2)Sulf(1)[S] | O-linked glycosylation | ② | ② |
| HNE+Delta_H(2)[K] | Chemical derivative | ② | ② |
| Malonyl[C] | Chemical derivative | ② | ② |
| MTSL[C] | Chemical derivative | ② | ② |
| Oxidation+NEM[C] | Chemical derivative | ② | ② |
| Phe->Thr[F] | AA substitution | ② | ② |
| PhosphoUridine[Y] | Post-translational | ② | ② |
| Ser->Trp[S] | AA substitution | ② | ② |
| SulfanilicAcid_13C(6)[E] | Chemical derivative | ② | ② |
| Xle->Phe[I] | AA substitution | ② | ② |
| Xlink_SMCC[219][C] | Chemical derivative | ② | ② |
| Xlink_SMCC[237][C] | Chemical derivative | ② | ② |
| Thiophospho[S] | Other | 66.16 | 0.0010 |
| NHS-LC-Biotin[AnyN-term] | Chemical derivative | 58.75 | 0.0142 |
| Ub-Br2[C] | Chemical derivative | 19.24 | 0.0155 |
| 4-ONE[H] | Chemical derivative | 17.34 | 0.0251 |
| Lys->Pro[K] | AA substitution | 12.42 | 0.0436 |
| Xle->Thr[I] | AA substitution | 12.32 | 0.0436 |
| Acetyl[AnyN-term] | Multiple | 11.08 | 0.0007 |
| Oxidation[E] | Chemical derivative | 10.95 | 0.0314 |
| methylsulfonylethyl[C] | Chemical derivative | 10.75 | 0.0233 |
| O-Methylphosphate[S] | Chemical derivative | 9.42 | 0.0011 |

| | | | |
|---|---|---|---|
| SulfurDioxide[C] | Post-translational | 8.68 | 0.0452 |
| Phosphoadenosine[T] | Post-translational | 8.63 | 0.0271 |
| Carbamyl[AnyN-term] | Multiple | 8.04 | 0.0003 |
| Gly->His[G] | AA substitution | 7.96 | 0.0316 |
| Asn->Thr[N] | AA substitution | 7.95 | 0.0069 |
| Asn->Cys[N] | AA substitution | 7.37 | 0.0155 |
| Isopropylphospho[T] | Chemical derivative | 7.15 | 0.0018 |
| AccQTag[AnyN-term] | Chemical derivative | 7.01 | 0.0018 |
| Cys->SecNEM_2H(5)[C] | AA substitution | 6.97 | 0.0025 |
| Carboxymethyl[K] | Chemical derivative | 6.82 | 0.0000 |
| NEMsulfur[C] | Chemical derivative | 6.67 | 0.0012 |
| Ethyl+Deamidated[Q] | Chemical derivative | 6.58 | 0.0313 |
| Crotonaldehyde[C] | Other | 6.17 | 0.0019 |
| Hydroxymethyl[N] | Post-translational | 5.83 | 0.0148 |
| Gly->Pro[G] | AA substitution | 5.59 | 0.0005 |
| Acetyl_2H(3)[AnyN-term] | Artefact | 5.48 | 0.0284 |
| Carboxyethyl[K] | Post-translational | 5.45 | 0.0249 |
| Oxidation[Y] | Post-translational | 5.36 | 0.0477 |
| Cyano[C] | Post-translational | 5.33 | 0.0035 |
| Piperidine[AnyN-term] | Chemical derivative | 5.32 | 0.0481 |
| Cation_Ni[II][D] | Artefact | 5.27 | 0.0354 |
| Nmethylmaleimide+water[C] | Chemical derivative | 5.25 | 0.0000 |
| MercaptoEthanol[T] | Chemical derivative | 5.20 | 0.0103 |
| Carbonyl[I] | Chemical derivative | 4.87 | 0.0075 |
| Val->Trp[V] | AA substitution | 4.68 | 0.0061 |
| Xle->Trp[I] | AA substitution | 4.58 | 0.0032 |
| Propionyl[AnyN-term] | Artefact | 4.14 | 0.0040 |
| Amidine[AnyN-term] | Chemical derivative | 4.11 | 0.0176 |
| DNCB_hapten[C] | Chemical derivative | 4.01 | 0.0002 |
| Phenylisocyanate_2H(5)[AnyN-term] | Chemical derivative | 3.65 | 0.0135 |
| dHex[N] | N-linked glycosylation | 3.51 | 0.0000 |
| C+12[AnyN-term] | * | 3.48 | 0.0092 |
| GIST-Quat_2H(3)[AnyN-term] | Artefact | 3.47 | 0.0118 |
| Asn->Trp[N] | AA substitution | 3.46 | 0.0347 |
| Trioxidation[C] | Chemical derivative | 3.42 | 0.0058 |
| Dioxidation[M] | Post-translational | 3.20 | 0.0198 |
| Unknown_210[AnyN-term] | Artefact | 3.18 | 0.0000 |
| MG-H1[R](Delta_H(2)C(3)O(1)[R]) | Other | 3.15 | 0.0011 |
| Tris[N] | Artefact | 3.08 | 0.0275 |
| Asn->Met[N] | AA substitution | 3.05 | 0.0452 |
| NEMsulfurWater[C] | Chemical derivative | 3.05 | 0.0007 |
| CarboxymethylDTT[C] | Artefact | 3.03 | 0.0006 |
| Cation_Fe[II][D] | Artefact | 3.02 | 0.0075 |
| Carboxy[W] | Chemical derivative | 3.01 | 0.0245 |

| | | | |
|---|---|---|---|
| Gly->Glu[G] | AA substitution | 2.87 | 0.0142 |
| Arg[AnyN-term] | Other | 2.75 | 0.0165 |
| Nmethylmaleimide[C] | Chemical derivative | 2.74 | 0.0020 |
| Oxidation[H] | Artefact | 2.65 | 0.0134 |
| Cation_K[D] | Artefact | 2.50 | 0.0354 |
| Cation_Fe[II][E] | Artefact | 2.36 | 0.0403 |
| Formylasparagine[H] | Chemical derivative | 2.35 | 0.0472 |
| Xle->Gln[I] | AA substitution | 2.35 | 0.0084 |
| Delta_H(2)C(2)[AnyN-term] | Other | 2.34 | 0.0309 |
| Cysteinyl[C] | Multiple | 2.30 | 0.0237 |
| Biotin_Thermo-88317[Y] | Chemical derivative | 1.96 | 0.0178 |
| Formyl[AnyN-term] | Artefact | 1.77 | 0.0105 |
| Xlink_EGS[115][K] | Chemical derivative | 1.76 | 0.0190 |
| Oxidation[W] | Artefact | 1.73 | 0.0164 |
| Cys->Dha[C] | AA substitution | 1.65 | 0.0066 |
| Gln->pyro-Glu[AnyN-termQ] | AA substitution | 0.34 | 0.0401 |
| dichlorination[Y] | Artefact | 0.29 | 0.0223 |
| azole[C] | Post-translational | 0.12 | 0.0037 |

Table S5 Details of differential modifications in two comparisons by open search.

* Modification type not retrieved in UNIMOD database;

① The unique modification type of the EW0 group; ② The unique modification types of the EM5 group; ③ The unique modification types of the CM5 group.

| Group | Modification Name | Modification Type | P-value | Fold Change | References |
|---|---|---|---|---|---|
| EM5 - EW0 | Carbamyl[AnyN-term] | Multiple | 0.0000 | 4.65 | [136] |
| | CHDH[D] | Post-translational | 0.0005 | 2.31 | — |
| | Trp->Kynurenin[W] | Chemical derivative | 0.0375 | 2.04 | [137] |
| | NHS-LC-Biotin[AnyN-term] | Chemical derivative | 0.1206 | 1.92 | — |
| | HNE+Delta_H(2)[K] | Chemical derivative | 0.2029 | 1.69 | [142] |
| | Thiazolidine[W] | Chemical derivative | 0.9862 | 1.00 | [145] |
| | Carboxyethyl[K] | Post-translational | 0.7766 | 0.91 | [146] |
| | PhosphoUridine[Y] | Post-translational | 0.7787 | 0.90 | — |
| | glucosone[R] | Other | 0.6874 | 0.86 | [143] |
| | Hep[T] | O-linked glycosylation | 0.5316 | 0.83 | — |
| | Ethyl+Deamidated[N] | Chemical derivative | 0.1841 | 0.83 | — |
| | dHex[N] | N-linked glycosylation | 0.3045 | 0.74 | [147] |
| | AccQTag[AnyN-term] | Chemical derivative | 0.0364 | 0.70 | — |
| | 4-ONE[H] | Chemical derivative | 0.2781 | 0.52 | [143] |

| | | | | | |
|---|---|---|---|---|---|
| | Oxidation[P] | Post-translational | 0.0000 | 0.47 | [138] |
| | Cysteinyl[C] | Multiple | 0.0234 | 0.43 | [139] |
| | BITC[AnyN-term] | Chemical derivative | 0.1039 | 0.39 | [144] |
| | SulfurDioxide[C] | Post-translational | 0.0050 | 0.33 | [140] |
| | NO_SMX_SIMD[C] | Chemical derivative | 0.0003 | 0.31 | — |
| | Delta_H(2)C(3)[K] | Other | 0.0045 | 0.29 | [141] |
| | Guanidinyl[K] | Chemical derivative | 0.0093 | 2.72 | — |
| | Carboxymethyl[K] | Chemical derivative | 0.1476 | 2.46 | [151] |
| | PhosphoUridine[Y] | Post-translational | 0.0204 | 1.88 | — |
| | Carbamyl[AnyN-term] | Multiple | 0.0061 | 1.76 | [136] |
| | DeStreak[C] | Chemical derivative | 0.2293 | 1.33 | — |
| | glucosone[R] | Other | 0.5228 | 1.32 | [143] |
| | Crotonaldehyde[C] | Other | 0.4124 | 1.30 | [140] |
| | BITC[K] | Chemical derivative | 0.5487 | 1.22 | [144] |
| | Arg->Trp[R] | AA substitution | 0.4018 | 1.21 | — |
| | 2-hydroxyisobutyrylation[K] | Post-translational | 0.6458 | 1.15 | [150] |
| | Cysteinyl[C] | Multiple | 0.8368 | 1.09 | [139] |
| EM5 | 4-ONE[H] | Chemical derivative | 0.8445 | 1.06 | [142] |
| - | dHex[N] | N-linked glycosylation | 0.8163 | 1.05 | [147] |
| CM5 | phenylsulfonylethyl[C] | Chemical derivative | 0.9917 | 1.00 | — |
| | Delta_H(2)C(3)[K] | Other | 0.5922 | 0.93 | [141] |
| | Oxidation[Y] | Post-translational | 0.2648 | 0.86 | [152] |
| | SulfurDioxide[C] | Post-translational | 0.3743 | 0.80 | [140] |
| | Malonyl[C] | Chemical derivative | 0.3600 | 0.77 | [150] |
| | HNE+Delta_H(2)[K] | Chemical derivative | 0.5489 | 0.76 | [142] |
| | MG-H1[R](Delta_H(2)C(3)O(1)[R]) | Other | 0.2428 | 0.72 | [149] |
| | His->Ser[H] | AA substitution | 0.1070 | 0.60 | — |
| | Carboxyethyl[K] | Post-translational | 0.2808 | 0.53 | [146] |
| | Oxidation[P] | Post-translational | 0.1375 | 0.47 | [138] |
| | Delta_H(2)C(2)[AnyN-term] | Other | 0.0260 | 0.39 | [148] |
| | Dihydroxyimidazolidine[R] | Multiple | 0.0147 | 0.31 | — |

Table S6 Results of limited search of modifications
